# Extensive Intratumor Proteogenomic Heterogeneity Revealed by Multiregion Sampling in High-Grade Serous Ovarian Tumor Specimens

**DOI:** 10.1101/761155

**Authors:** Allison L. Hunt, Nicholas W. Bateman, Waleed Barakat, Sasha Makohon-Moore, Brian L. Hood, Kelly A. Conrads, Ming Zhou, Valerie Calvert, Mariaelena Pierobon, Jeremy Loffredo, Tracy J. Litzi, Julie Oliver, Dave Mitchell, Glenn Gist, Christine Rojas, Brian Blanton, Emma L. Robinson, Kunle Odunsi, Anil K. Sood, Yovanni Casablanca, Kathleen M. Darcy, Craig D. Shriver, Emanuel F. Petricoin, Uma N. M. Rao, G. Larry Maxwell, Thomas P. Conrads

## Abstract

Enriched tumor epithelium, tumor-associated stroma, and whole tissue were collected by laser microdissection from thin sections across spatially separated levels of ten primary high-grade serous ovarian tumors and analyzed using proteomics (mass spectrometry and reverse phase protein microarray) and RNA-sequencing analyses. Comparative analyses of transcript and protein abundances revealed independent clustering of enriched stroma and enriched tumor epithelium, with whole tumor tissue clustering between purified collections, driven by overall tumor purity. Comparison of historic prognostic molecular subtypes for HGSOC revealed protein and transcript expression from tumor epithelium correlated most strongly with the differentiated molecular subtype, whereas stromal proteins and transcripts most strongly correlated with mesenchymal subtype. Protein and transcript abundance in tumor epithelium and stromal collections from neighboring sections exhibited decreased correlation in samples collected just hundreds of microns apart. These data reveal substantial protein and transcript expression heterogeneity within the tumor microenvironment that directly bears on prognostic signatures and underscore the need to enrich cellular subpopulations for expression profiling.

## Introduction

Ovarian cancer is the fifth leading cause of death in women in the U.S. [1], with nearly 22,000 new cases and 14,000 deaths projected to occur in 2020. Most ovarian cancer cases are diagnosed at an advanced stage and exhibit heterogeneous populations and subpopulations of tumor cells that limit successful therapeutic intervention. The 5-year survival rate of patients diagnosed with metastatic ovarian cancer is less than 30% [2].

Numerous high-throughput sequencing studies aimed at broadly characterizing the genomic landscape of specific cancer types have been or are being conducted [3]. Although adding substantially to our molecular understanding of cancer, these studies have resulted in limited clinical translation, due in part to an ever increasing body of evidence that points to previously underappreciated levels of heterogeneity in the tumor microenvironment (TME) [4].

Substantial molecular and pathologic intra-tumoral differences in HGSOC tumors and their relationship with TME dynamics have been described [5, 6]. One study demonstrated that stable and/or regressing tumors lacked common neoepitopes and mutations compared to progressing tumors in the same patient [5], implicating non-somatic factors within the TME as critical determinants of immune response and overall tumor fate. Multiregional sampling has revealed extensive variation between subpopulations of cells within a single tumor [7–9], suggesting individual tumor samples have multiple subtype signatures present with differing levels of activation [10]. Single-cell RNA-seq of HGSOC samples has revealed grade-specific and cell type-specific transcriptional profiles representative of discrete cell types, such as tumor epithelium, stromal, and immune cells present within individual tumor specimens [7]. Further, the presence of subclonal cell populations within primary and/or metastatic tumors has been demonstrated to influence the state of immune infiltration and activation [8].

In this study, we investigated proteomic and transcriptomic heterogeneity in the TME from HGSOC patient primary tumors using laser microdissection (LMD) of spatially separated tumor enriched regions, stromal cell populations, as well as whole tumor collections throughout distinct levels from each specimen block. These results reveal stark molecular heterogeneity in the HGSOC tumor microenvironment and underscore the need to account for compartmental heterogeneity in the TME in molecular profiling analyses of HGSOC.

## Results

### Proteogenomic analyses of discrete cellular populations within a single HGSOC tumor using locoregional multi-sampling

Consecutive thin sections (∼200) were generated from fresh-frozen primary HGSOC tumors (n=10 patients) to support multi-region sampling by LMD followed by integrated quantitative proteomic and transcriptomic analyses (Fig. 1A; Supplemental Fig. 1). All patients had advanced stage (stage III or IV) ovarian or tubo-ovarian disease (Supplemental Table 1). Primary specimens were obtained from ovarian and/or tubo-ovarian tissue and other pelvic masses, to include omentum. Some of the patients were chemotherapy-naïve (n=6) at the time of surgical resection, while others had received neoadjuvant chemotherapy (n=4). Representative hematoxylin and eosin (H&E) stained sections were reviewed by a board-certified pathologist (UNMR) to confirm histologic characteristics and tumor cellularity within each level (Supplemental Table 1).

**Figure 1.**
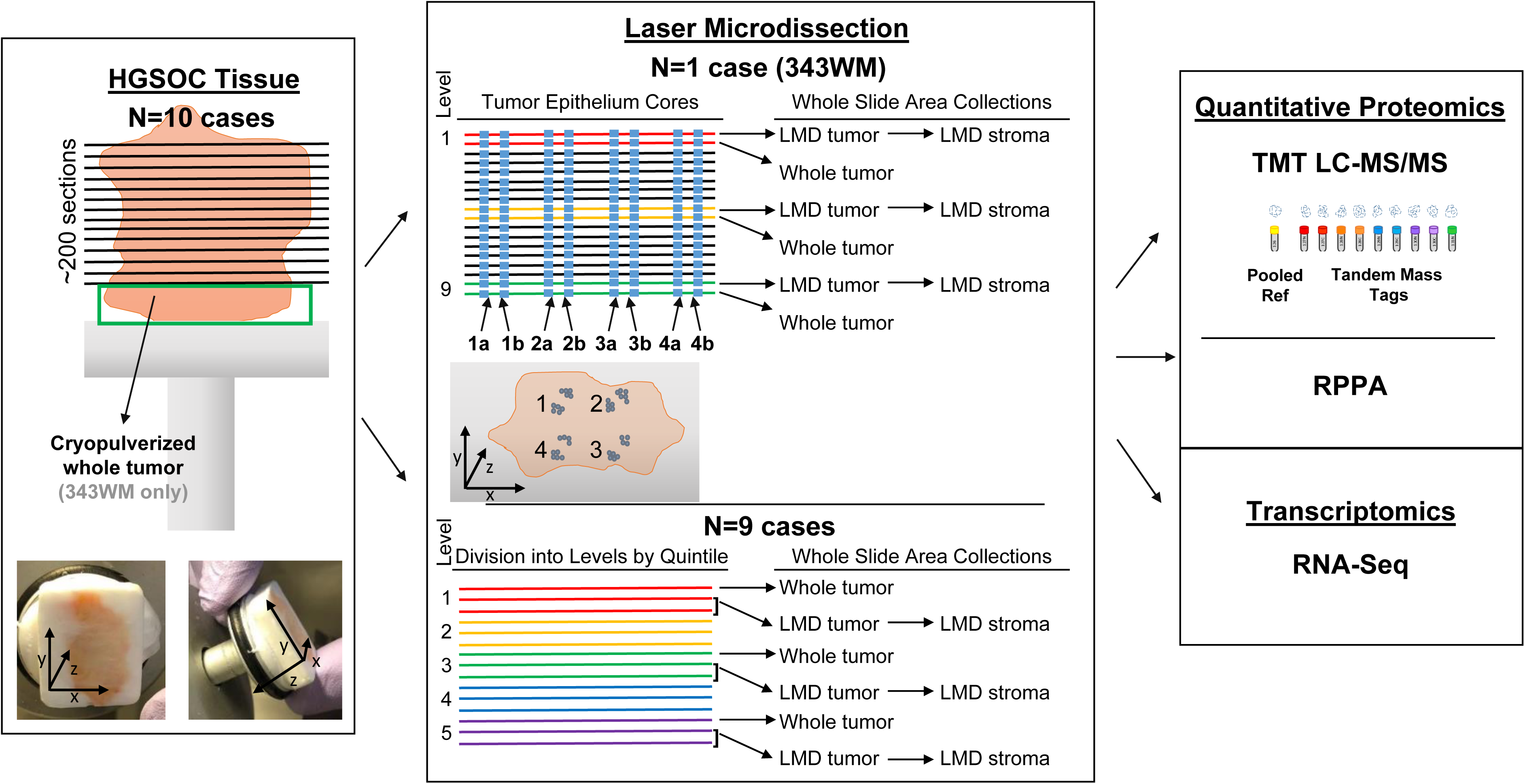
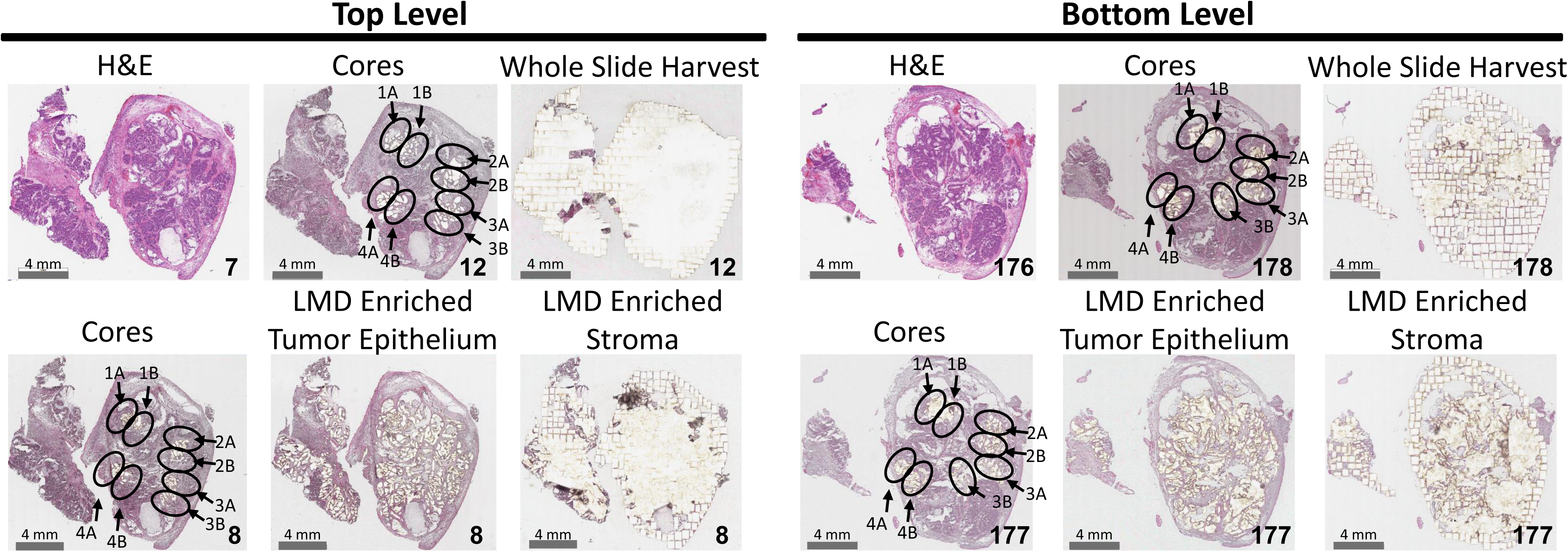
Study workflow. Illustration of histological tissue preparation, laser microdissection, proteomic analyses, and transcriptomic analysis (A), with representative pre- and post-LMD images from the top and bottom levels of the tissue from Case 343WM (B). (A) Specimen blocks obtained from 10 HGSOC patients were each cut into ∼200 consecutive thin tissue sections (left panel), which were each laser microdissected for enrichment of discrete tumor epithelium, discrete stroma, or whole tumor collections (middle panel) for analysis via quantitative proteomics and transcriptomics (right panel). One case (343WM; middle panel, top) was uniquely used for laser microdissection (LMD) enrichment of four tumor epithelium cores with adjacent replicate regions from each of 100 or 50 slides evenly distributed through the depth of the specimen for MS proteomics or transcriptomics, respectively. For 343WM, additional sets of 9 slides from spatially separated levels within the specimen block were each discretely microdissected for all remaining tumor and stroma after collecting the cores by LMD, as well as a nearest neighboring whole tumor collection. The remainder of the specimen was cryopulverized in liquid nitrogen. The specimen blocks from the remaining 9 cases (middle panel, bottom) were divided into 5 levels (quintiles) of equal depth. Within each level, interlaced sections were used for LMD enrichment of discrete tumor epithelium, discrete stroma, and whole tumor collections for each downstream analytical purpose. Proteins and transcripts isolated from each of these distinct collections were analyzed by isobaric tagging and high-resolution liquid chromatography-tandem mass spectrometry, reverse phase protein microarray, and/or next generation sequencing. (B) Representative pre- and post-LMD images are shown for 343WM from tissue sections at the top and bottom levels used for proteomic analysis. The number in the bottom right corner of each micrograph indicates the slide number shown. The scale bar in the bottom left corner of each micrograph indicates a length of 4 mm.

Enriched tumor epithelium, stromal cells, as well as whole tumor collections representing all material on a single tissue section, were collected at spatially distinct intervals (“levels”) from alternating sections throughout the tumor blocks (Fig. 1B; Supplemental Table 2). In case 343WM, additional LMD collections were conducted to generate spatially separated “cores” (with additional technical replicate “cores”) representing defined sub-populations of tumor epithelium from alternating tissue sections that spanned the entirety of the patient tumor block from four spatially diverse quadrants (i.e., 1 - 4) and combined into four independent sample sets, including close biological collection “replicates” (i.e. 1A, 1B, 2A, 2B, etc) to support comparative proteome and transcriptome analyses (Fig. 1B). Representative PEN membrane slides after LMD were imaged, along with adjacent H&E stained tissue sections, to enable co-registration and automated quantification of tumor and stromal cell populations collected (Supplemental Table 3). There was marked discrepancy regarding estimation of tumor purity between the automated and manual evaluations. Digital analysis estimated 23 - 56% tumor cellularity at different depths of the block, calculated based on tumor regions of interest (ROI) collected by LMD / whole tissue area (Supplemental Table 3). Further, automated estimates revealed median cell number per area to be approximately 7,482 cells/mm^2^. By cell type, this represents an average of 7,527 ± 172 tumor cells (n=15 protein and RNA collections) or 3,899 ± 263 stroma cells (n=6 protein and RNA collections) collected by LMD per mm^2^ of tissue area. Comparatively, manual review estimated 75 - 95% tumor cellularity (Supplemental Table 1). Differences between automated and manual review of tumor cellularity can likely be attributed to the presence of small regions of interceding stroma and other cell types in and around the tumor epithelium, which were excluded from the LMD tumor epithelium collections.

Multiregion protein samples were analyzed by mass spectrometry-based quantitative proteomics using a multiplexed, isobaric tagging methodology (tandem mass tags, TMT-10 or TMTpro-16; Supplemental Table 4) and reverse phase protein microarray (RPPA). RNA samples were analyzed by targeted RNA-seq analyses. For 343WM, a total of 6,053 proteins (Supplemental Table 5) and 20,535 transcripts (Supplemental Table 6) were quantified across all samples. For the remaining 9 cases, an average of 9,223 ± 511 proteins were measured within each patient-specific TMT plex for a total of 6,199 proteins co-quantified across all patients (Supplemental Table 7). A total of 19,758 transcripts were quantified from 343WB and 343WH (Supplemental Table 8).

### Tumor and stromal cell populations exhibit diverse proteogenomic profiles

Unsupervised hierarchical cluster analysis of differentially expressed proteins (Supplemental Table 9) or transcripts (Supplemental Table 10) from case 343WM revealed distinct sub-clustering of tumor cores, enriched tumor epithelium and stroma, as well as whole tumor collections (Fig. 2A and B). Two predominant clusters stratifying tumor cores and tumor epithelium from stroma and whole tumor collections are apparent. Notably, the cryopulverized tissue proteome exhibited an intermediate cluster between these two sample groups (Fig. 2A), suggesting this sample type represents a mixture of these cellular populations. The proteins measured in the cryopulverized tissue also reflect contributions from the inclusion of blood, necrosis, and fat within this sample, which had been excluded from the whole tumor collections.

**Figure 2.**
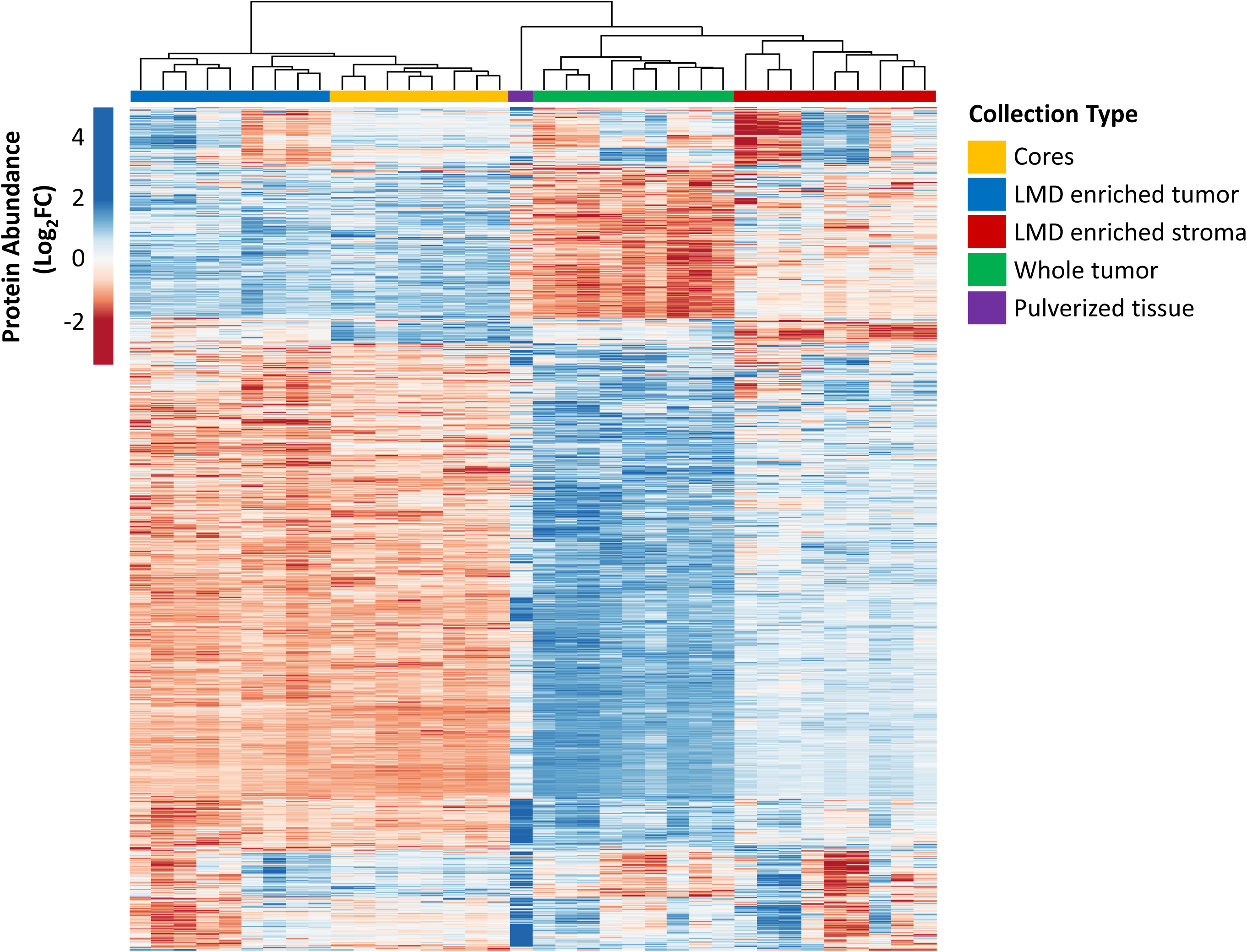
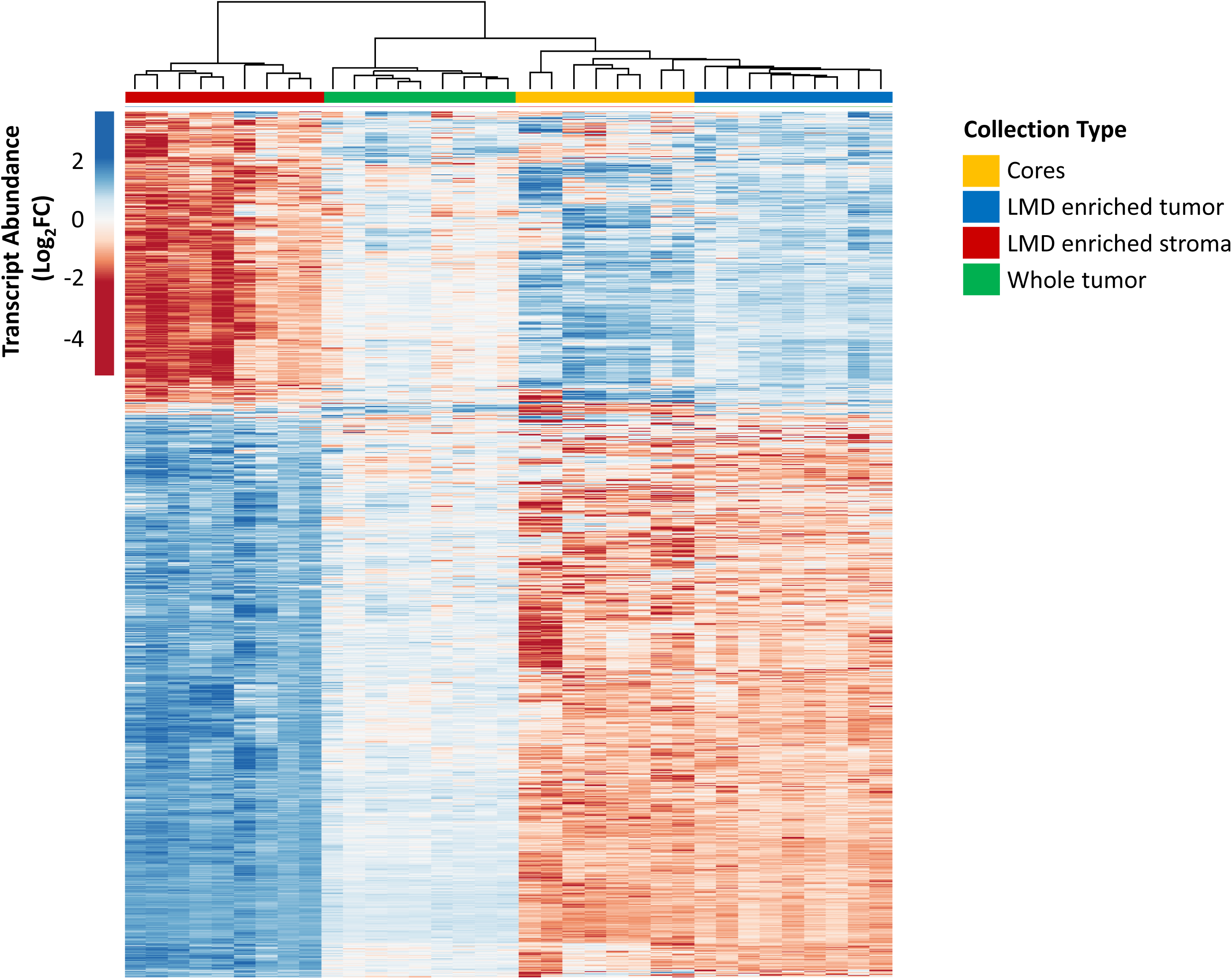
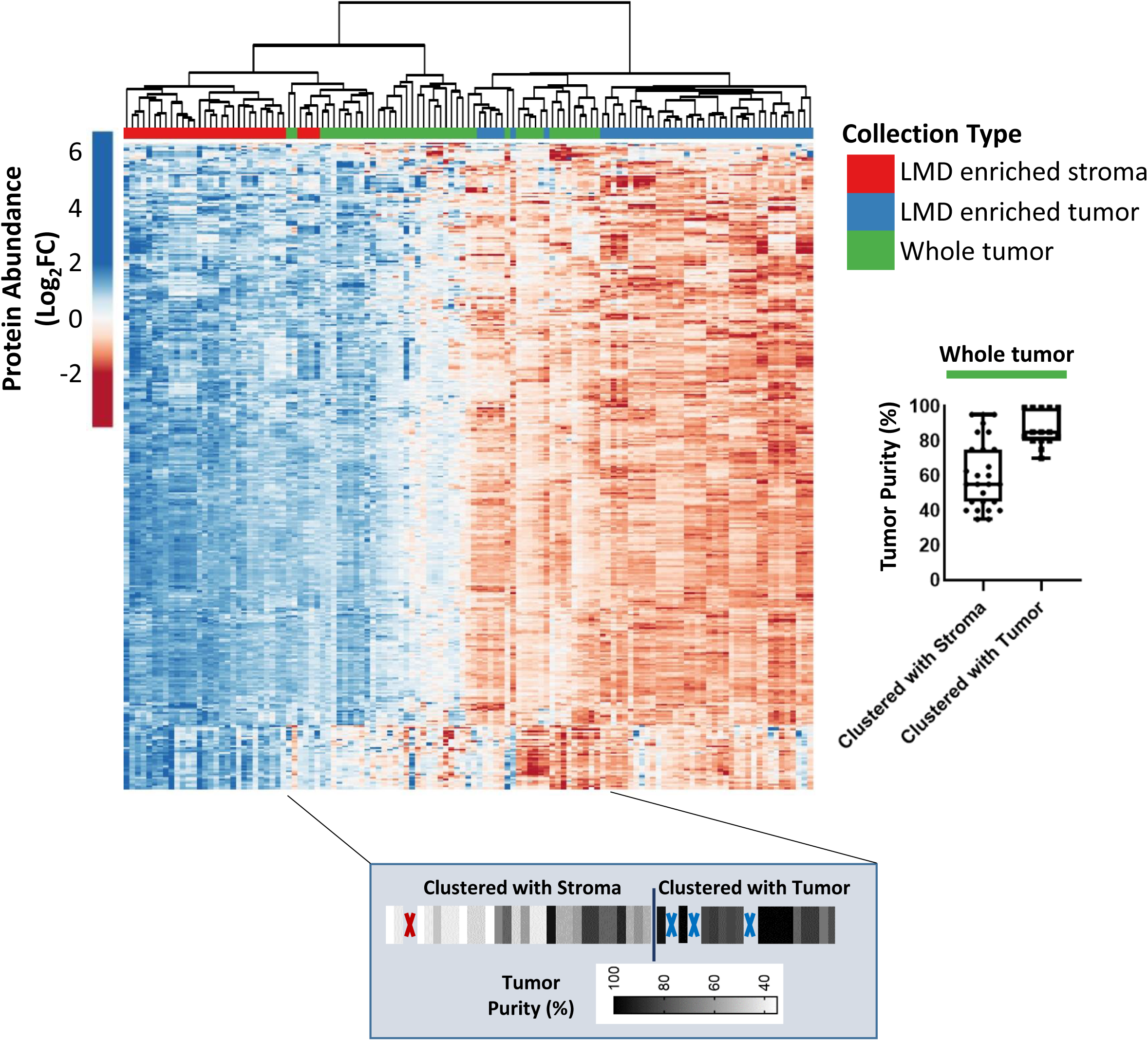
Unsupervised hierarchical cluster analysis of differentially abundant proteins and transcripts. (A) 1,928 differentially abundant proteins and (B) 3,861 transcripts with median absolute deviation (MAD)>0.5 from case 343WM, and (C) 6,199 differentially abundant proteins with MAD>1 in the expanded cohort (n=9). (C) Protein abundances are represented across n=123 samples derived from n=9 patients in the expanded cohort consisting of: LMD enriched tumor epithelium (n=45 total; 5 levels/patient), LMD enriched stroma (n=33 total; 2-5 levels/patient), and whole tumor (n=45 total; 5 levels/patient) samples. Highlighted box depicts the median tumor purity values from manual pathology review for each of the whole tumor collections as they appear in the heatmap, ordered from left to right. Dark elongated border line distinguishes the whole tumor collections which clustered with LMD enriched stroma (left) from those that clustered with LMD enriched tumor (right). Red and blue “X” marks represent interceding LMD enriched stroma or tumor collections, respectively.

Similar clustering was seen across the additional 9 HGSOC patient tissue cell type collections; unsupervised hierarchical cluster analysis of proteins (Supplemental Table 11) and transcripts (Supplemental Table 12) revealed distinct clustering of the LMD enriched tumor epithelium from the LMD enriched stroma (Fig. 2C; Supplemental Fig. 2). Importantly, the extent to which the whole tumor tissue collection more closely associated with tumor epithelium or stroma was directly related to the tumor “purity” reflecting levels of overall tumor purity; namely, whole tumor tissues with high percent tumor epithelium associated with the LMD enriched tumor epithelium and those with low percent tumor epithelium more strongly associated with LMD enriched stroma (Supplemental Table 1).

We next examined the abundance of representative epithelial markers Ovarian Cancer-Related Tumor Marker CA125/Mucin-16 (CA-125/*MUC16*), Keratin Type I Cytoskeletal 19 (*KRT19*), and Cadherin 1 (*CDH1*), and stromal markers Versican (VCAN) and Fibroblast Activation Protein Alpha (FAP) across each tissue collection type (Fig. 3A). As expected, MUC16 and CDH1 protein and transcript levels were more abundant in tumor cores and LMD enriched tumor epithelial collections, and comparatively decreased in LMD enriched stroma and whole tumor collections. VCAN and FAP abundance, in contrast, were elevated in LMD enriched stroma and whole tumor collections, consistent with these proteins being stromal in origin [11, 12]. Cell type enrichment analyses using transcript expression-level data from 343WM was performed (Supplemental Table 13) [13], and except for Cores 4A/B, all of the LMD tumor cores and LMD enriched tumor epithelium collections correlated most strongly with epithelial cell signatures, while the LMD enriched stroma and whole tumor collections had the highest correlation with fibroblast cell signatures, stroma, as well as microenvironment scores (Fig. 3A). Overall, these highly significant (p<0.0001) associations between epithelial and stromal marker proteins KRT19, CDH1, VCAN, and FAP with LMD enriched tumor epithelium and stroma, respectively, were observed in the additional 9 HGSOC cases assessed (Fig. 3B). Cellular type enrichment scores were also significantly different in the additional 9 cases, in which LMD enriched tumor was again significantly enriched for epithelial cells, while LMD enriched stroma had the highest scores for fibroblasts, stroma, and microenvironment (p<0.0001; Fig. 3C; Supplemental Table 14).

**Figure 3.**
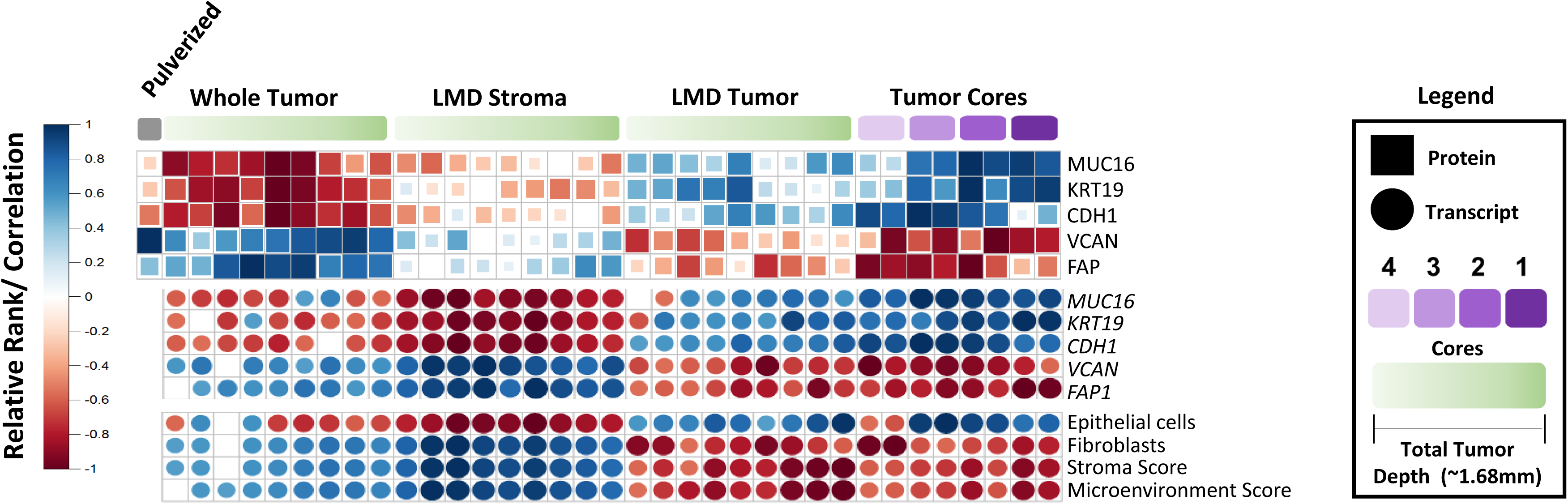
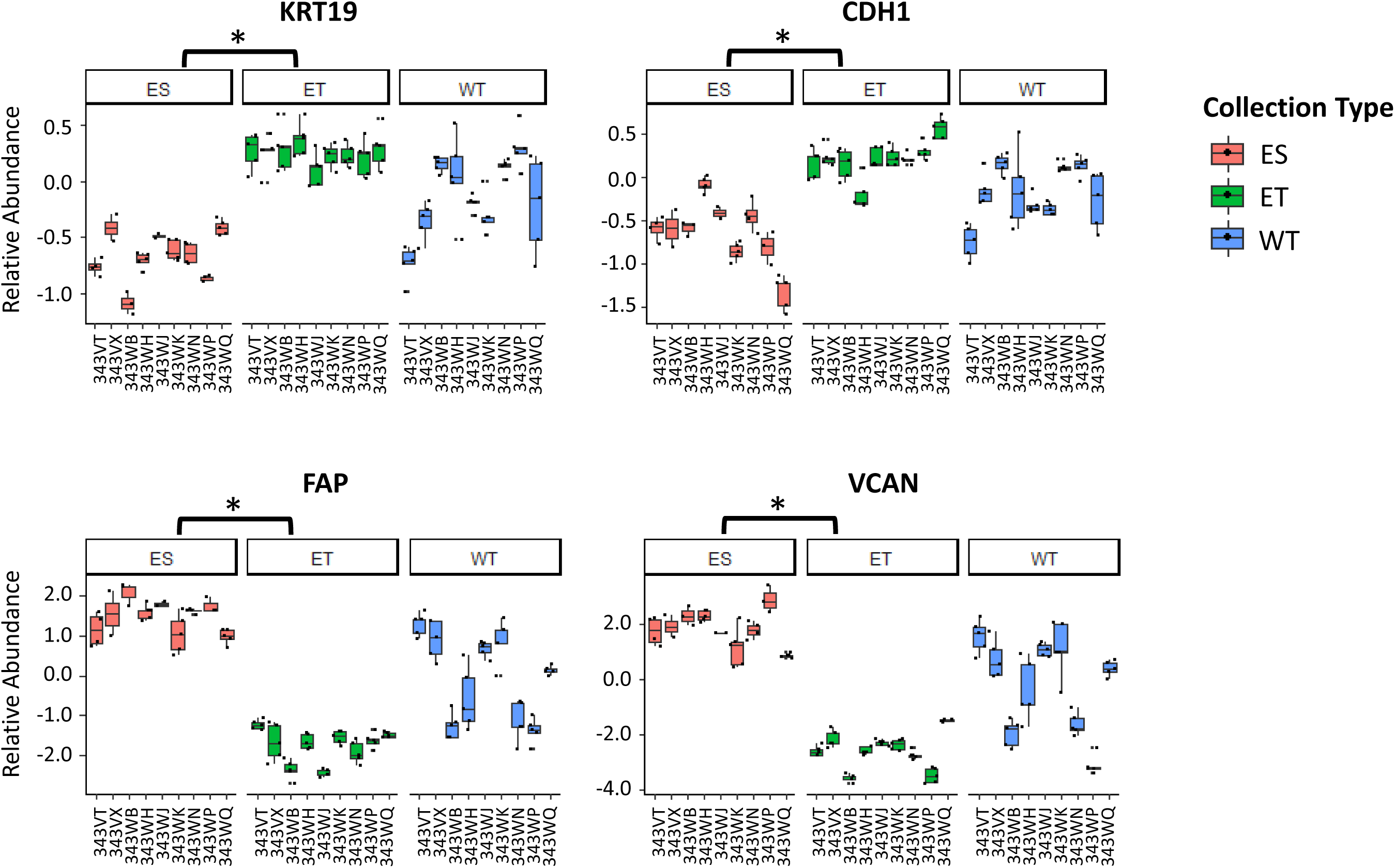
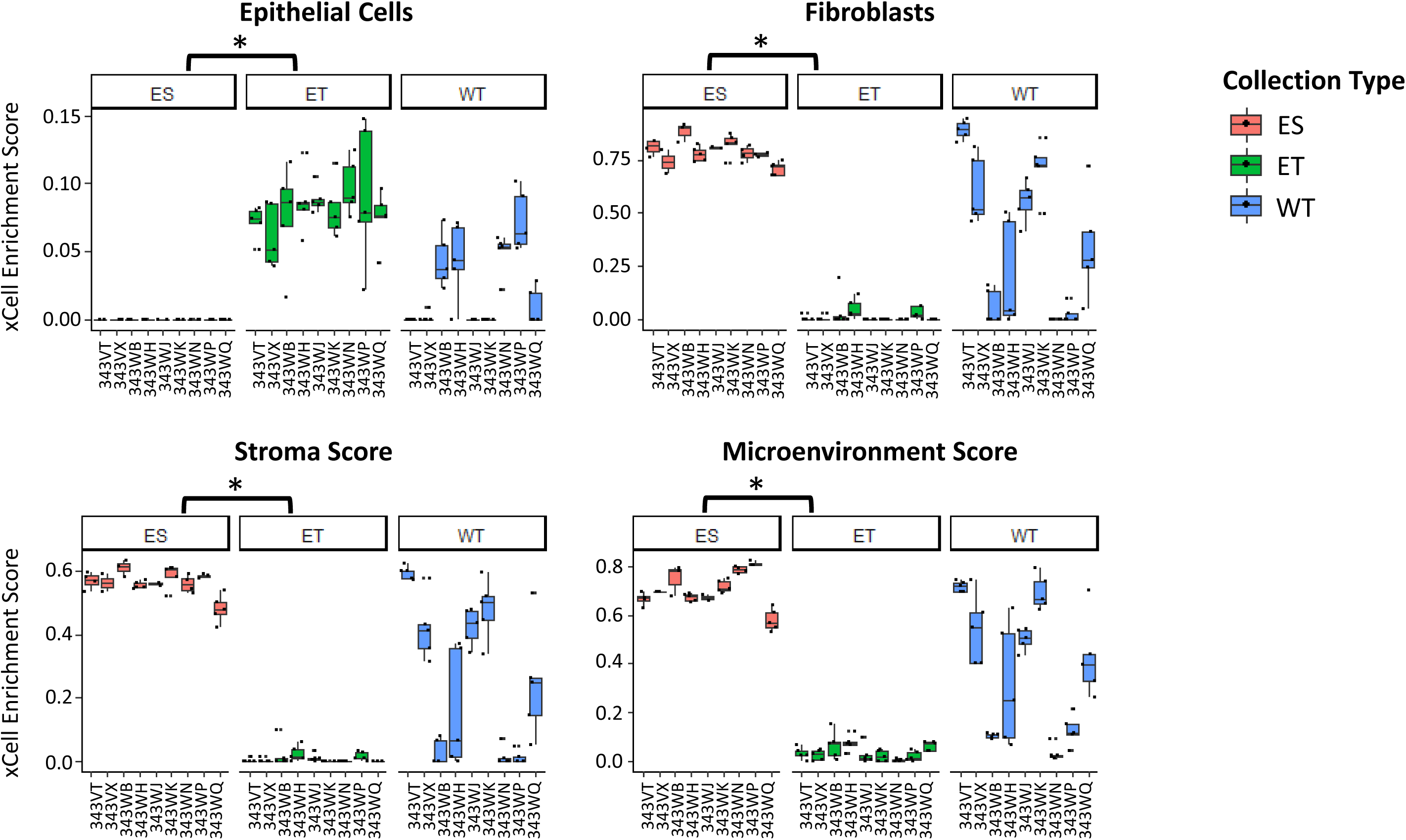
Protein and transcript abundance of epithelial and stromal markers in HGSOC as well as cellular admixture analyses (xCell. [13]). (A) Heatmaps depicting protein and transcript abundances and cellular admixture enrichment scores from 343WM for replicate tumor cores and by depth within the specimen block for LMD enriched tumor, stroma, and whole tumor collections. Protein abundance from the cryopulverized tumor is included. Size and color of each shape reflects Spearman correlation. (B) Boxplots depicting relative protein abundances for KRT19, CDH1, FAP, and VCAN and (C) cellular admixture scores in the expanded cohort (n=9). ES= LMD enriched stroma; ET= LMD enriched tumor; WT= whole tumor. P-values with (*) indicate statistically significant differential expression (p<0.0001) between ES and ET.

Notably, the abundance of MUC16 was not significantly different between LMD enriched tumor and stroma (p=0.0825; Supplemental Fig. 3) in the proteomic datasets of the additional 9 patients. MUC16 was elevated in LMD enriched stroma for cases 343VT and 343WQ and was not measured in any tissue collections from cases 343WH and 343WN. CA-125 is monitored as a clinical biomarker in serum for ovarian cancer progression and regression [14–16] and is an epitope of the extracellular domain of the MUC16 protein [17–19] which can be shed from HGSOC cells via proteolytic cleavage [20]. As MUC16 was not significantly altered between LMD enriched tumor and stroma, we hypothesized the abundance of proteins shed or secreted from tumor cells may exhibit greater variability within the tumor microenvironment due to circulation within interstitial spaces as well as local vasculature. We therefore compared abundance trends of proteins exhibiting a signal peptide sequence for targeting a protein for secretion to the extracellular space, or that are characterized as being extracellular by Gene Ontology, relative to proteins lacking these features across all collection levels to measure whether the variance in protein abundance across levels was higher in proteins with versus without signal peptides or extracellular localization. Cases 343WM, 343WK, 343WQ, and 343WH were selected for this analysis as proteomic data from all levels/case of each LMD enriched tumor and stroma was available. We found that proteins with signal peptide sequences expressed in the LMD enriched tumor epithelium in three cases and proteins in the LMD enriched stroma from all four cases had significantly higher variance (p<0.0001) than proteins lacking these sequences (Supplemental Table 15). Further, the variance of extracellular proteins was also significantly higher (p<0.001) than those lacking this classification.

Integrated analyses of protein and transcript level data revealed 5,742 co-quantified proteins and transcripts from 343WM across all tissue types (Fig. 4; Supplemental Table 16). Proteins and their transcripts for a given LMD collection type showed positive correlation with other samples from the same LMD collection type (e.g. Core 1A protein clustered most strongly to its corresponding transcript and next nearest to other core collections). Notably, stromal transcripts clustered most strongly with their cognate proteins from the whole tumor collections. In comparison of replicate tumor cores, the strongest correlations, apart from intra-core comparisons, were between the “A” and “B” replicates for each core. The A and B replicates for each core at the proteome-only and transcriptome-only levels had Spearman Rho between 0.65 - 0.91 and 0.75 - 0.84, respectively. For replicate cores at the protein and corresponding transcript levels the Spearman Rho were 0.38 - 0.51. In analyses of protein and transcript abundance in correlations in other patient tumors, there were additionally 5,379 co-quantified proteins and transcripts measured from LMD enriched tumor, stroma, and whole tumor from one level each for cases 343WH and 343WB (Supplemental Table 17). Correlations were strongest within the same LMD collection type (Supplemental Figure 2). For 343WH, the LMD enriched stroma proteins correlated most strongly with the corresponding LMD enriched stroma transcripts, but was also positively correlated with whole tumor transcripts, consistent with 343WM. The tumor cellularity was marginally higher in 343WH (median tumor cellularity 85 - 95%) relative to 343WM (median tumor cellularity 80 - 95%) (Supplemental Table 1) which likely accounts for differences in relatedness between the LMD enriched stroma protein and either whole tumor RNA or LMD enriched stroma RNA,

**Figure 4.**
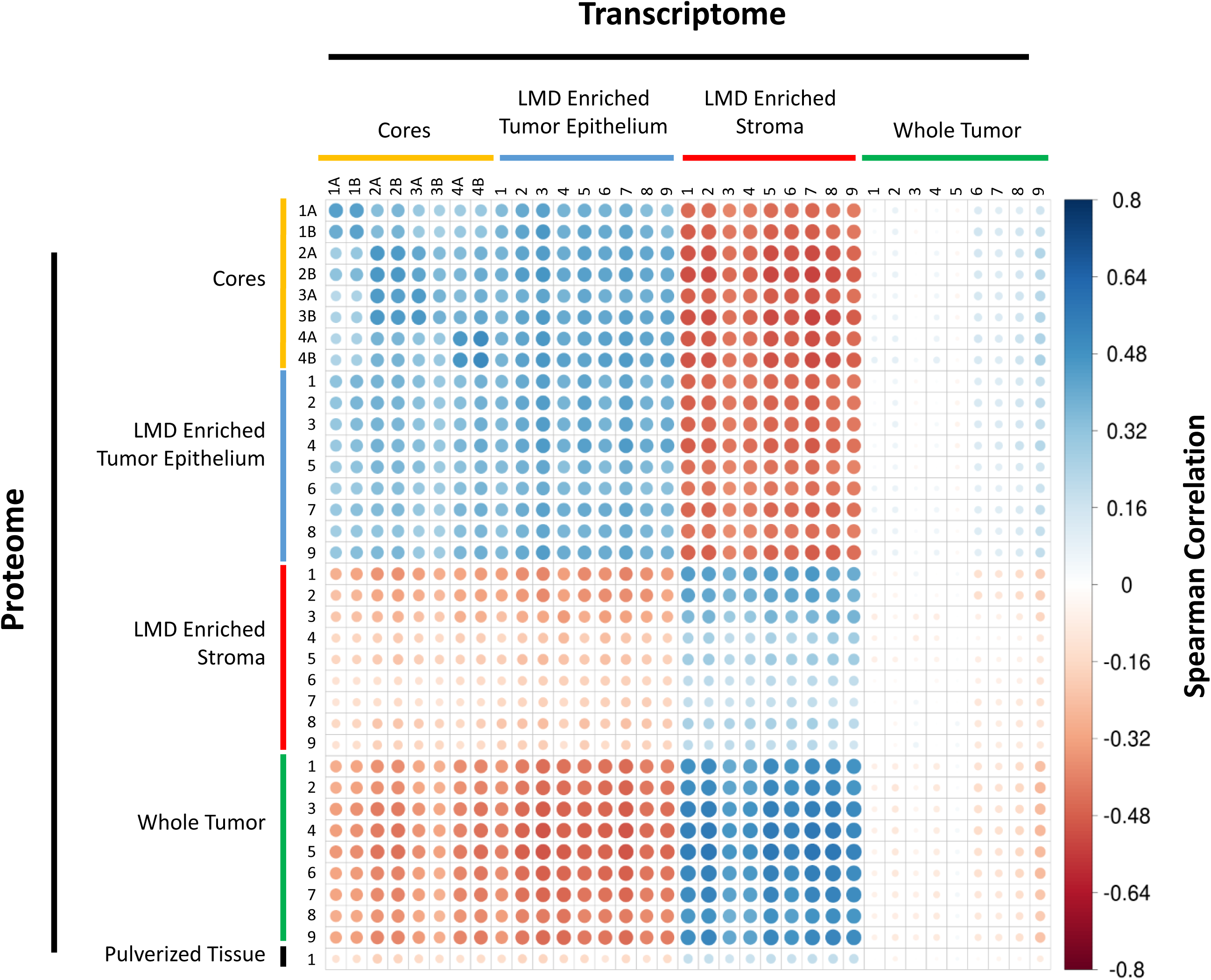
Protein-RNA Spearman Correlation Matrix for case 343WM. Spearman correlation analysis of 5,742 genes that were co-measured as proteins and corresponding transcripts in 343WM. Size and color of each circle reflects Spearman correlation.

We further used 281 antibodies (Supplemental Table 18) against native and/or post-translationally modified proteins for RPPA analysis (Supplemental Table 19). Integrated analyses of MS-based and RPPA proteome data (Supplemental Table 20) prioritized 54 proteins as co-quantified between platforms and revealed similar findings as above indicating generally positive correlation of protein abundances between LMD enriched tumor (Spearman Rho between -0.24 and 0.59; Supplemental Fig. 5A) and stroma (Spearman Rho between 0.09 and 0.61; Supplemental Fig. 5B) across all cases, with the exceptions of LMD enriched tumor from 343VT and, to a lesser extent, 343WH showing mixed relatedness to other cases. However, the whole tumor collections (Supplemental Fig. 5C) displayed markedly poorer concordance (Spearman Rho between -0.50 and 0.49) between the two analytical platforms.

**Figure 5.**
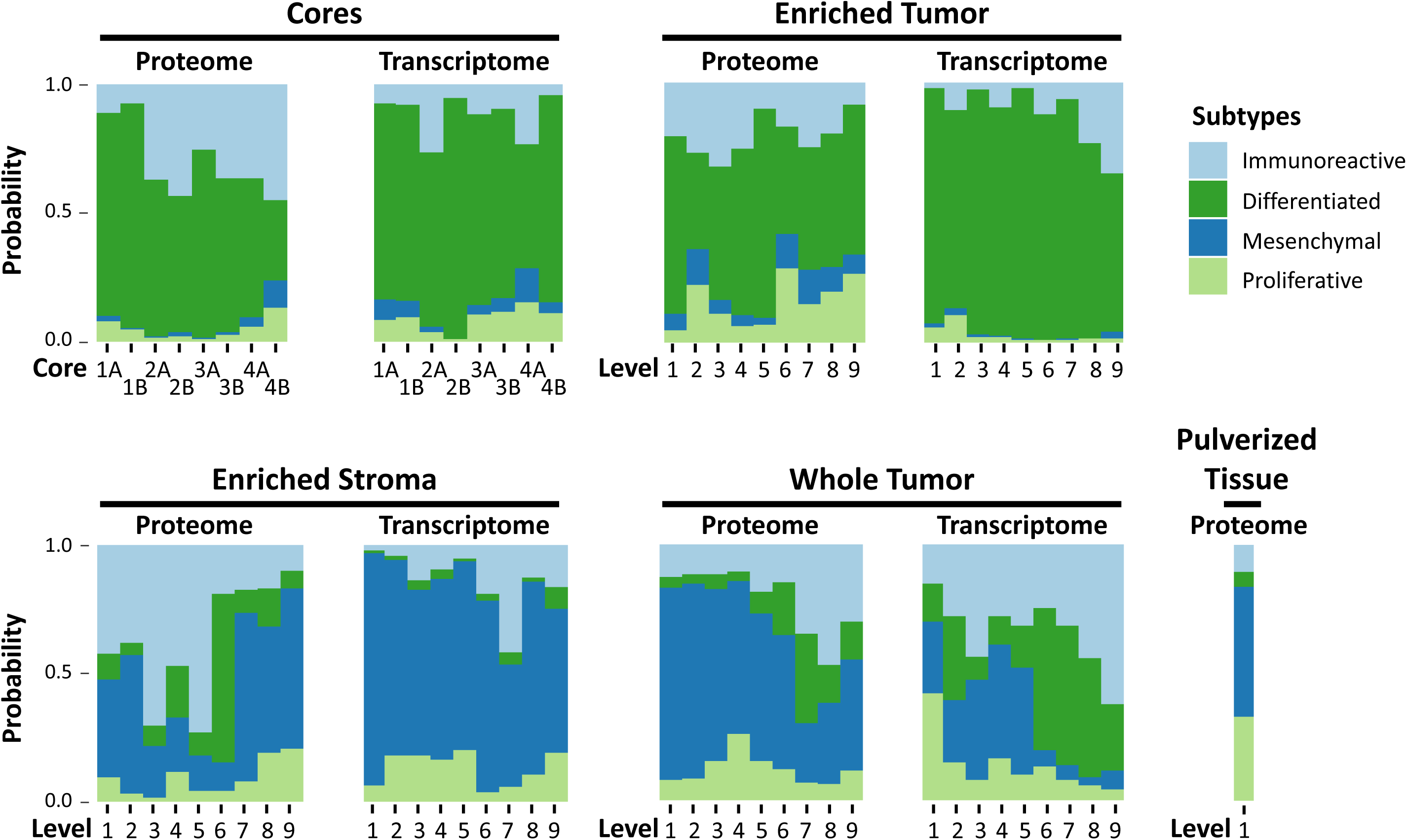
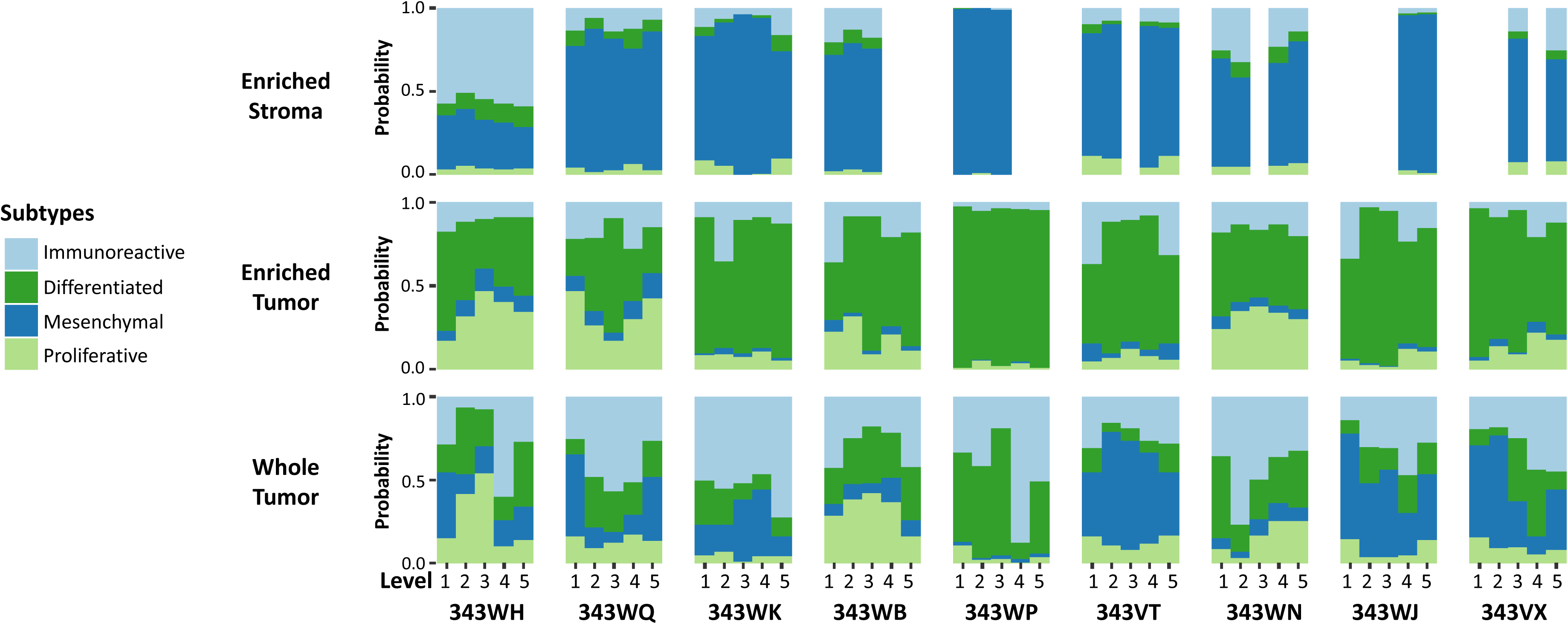
Protein and transcript abundance of markers correlating with prognostic molecular signatures of altered disease outcome in HGSOC. (A) Transcript abundances for markers correlated with suboptimal surgical debulking identified by Liu et al (2015) [24], and protein and transcript abundances correlating with prognostic molecular subtypes identified by Konecny et al (2014) [22] measured in 343WM. (B) Stacked bar graphs depicting the probability of each collection type (LMD enriched stroma, LMD enriched tumor, and whole tumor) per level per patient belonging to consensus molecular subtypes identified by Chen et al (2018) [4] based on protein abundance.

Comparison of proteomic and transcriptomic data revealed common canonical pathways (Ingenuity Pathway Analysis, IPA) that were consistently altered within LMD collection types (Supplemental Table 21). Aryl hydrocarbon receptor signaling, and the endocannabinoid cancer inhibition pathway were among the top 5 canonical pathways most activated in LMD enriched tumor epithelium relative to stroma at both the proteomic and transcriptomic level. Conversely, IL-8 signaling and GP6 signaling were among the pathways predicted to be inhibited, i.e. least activated in enriched tumor epithelium relative to stroma. Additional pathways differentially expressed between tumor and stroma included activation of organismal death, morbidity and mortality, bleeding, organization of the cytoplasm and skeleton, and cell movement and migration in both proteome and transcriptome level data (Supplemental Table 22). Genes encoding known drug targets of FDA-approved drugs [21] which were differentially expressed (LIMMA adjusted p-value <0.01) between the LMD enriched tumor and LMD enriched stroma were identified in the proteomic and transcriptomic data with Log_2_ fold-changes > ±1 are listed in Supplemental Table 23. The FDA-approved drugs targeting the identified genes include a mixture of those being used currently for clinical management of HGSOC disease, such as bevacizumab, as well as many drugs that are not. Similar results were not seen using the proteomic data from case 343WM.

For the additional 9 HGSOC cases, we examined pathway enrichment using data for 381 proteins which displayed the same pattern of enrichment across all 9 cases, and which had Log_2_ fold-changes > ±1 (Supplemental Table 24). Overall, the canonical pathways (Supplemental Tables 25, 26) enriched in LMD tumor epithelium relative to stroma generally overlapped with those from case 343WM suggesting these pathways are highly enriched within tumor or stroma cell populations of HGSOC tumors. The proteomic data from these 9 cases revealed protein targets of olaparib, estramustine, sorafenib, regorafenib, and pazopanib that were differentially expressed between LMD enriched tumor epithelium and stroma (Supplemental Table 27).

### Intratumoral proteogenomic heterogeneity impacts prognostic molecular signatures correlating with altered disease outcome in HGSOC

Several large-scale studies have aimed to identify prognostic molecular signatures for HGSOC patient outcome, most of which have been from genomic and/or transcriptomic data from whole tumor specimen preparations from tumors that are typically qualified for inclusion using tumor cellularity/purity thresholds determined from pathologic inspection [3, 10, 22–26]. Per the categorization of tumors into immunoreactive, differentiated, proliferative, and mesenchymal subtypes, patients whose tumors are characterized by mesenchymal signatures have the worst prognosis. Consensus signatures generated by Chen *et al* [4] represent molecular subtype classifications and a random forest probability for composite classifiers generated from four historical HGSOC molecular subtypes classification systems [10, 22, 27, 28]. We used these consensus signatures to examine correlations between each spatially resolved tissue collection type to these HGSOC prognostic sub-types (Fig. 5; Supplemental Table 28; Supplemental Table 29). This analysis revealed that transcripts from LMD-enriched tumor cores and tumor epithelium from case 343WM were strongly correlated with the differentiated and/or proliferative subtypes and anti-correlated with the prognostically-poor mesenchymal subtype (Fig. 5A). Conversely, transcripts from the LMD-enriched stroma most strongly correlated to the prognostically-poor mesenchymal subtype. Molecular subtype profiles for enriched tumor and stroma were largely recapitulated at the proteome level for case 343WM (Fig. 5A) and in the additional 9 HGSOC cases (Fig. 5B). The whole tumor collections showed various levels of correlation to all molecular subtypes, the extent to which seemingly depends on the percent stromal and tumor epithelium admixture contributions to overall tumor purity.

We additionally investigated performance of a suboptimal cytoreduction associated network (SCAN) signature [24] correlating with stroma activation and tumor invasiveness in transcriptome data from case 343WM. The 11-gene surgical debulking signature from Liu *et al.* most strongly correlated with our LMD-enriched stroma and whole tumor tissue collections, (Supplemental Fig. 6), consistent with the correlation of the SCAN signature with stromal involvement.

**Figure 6.**
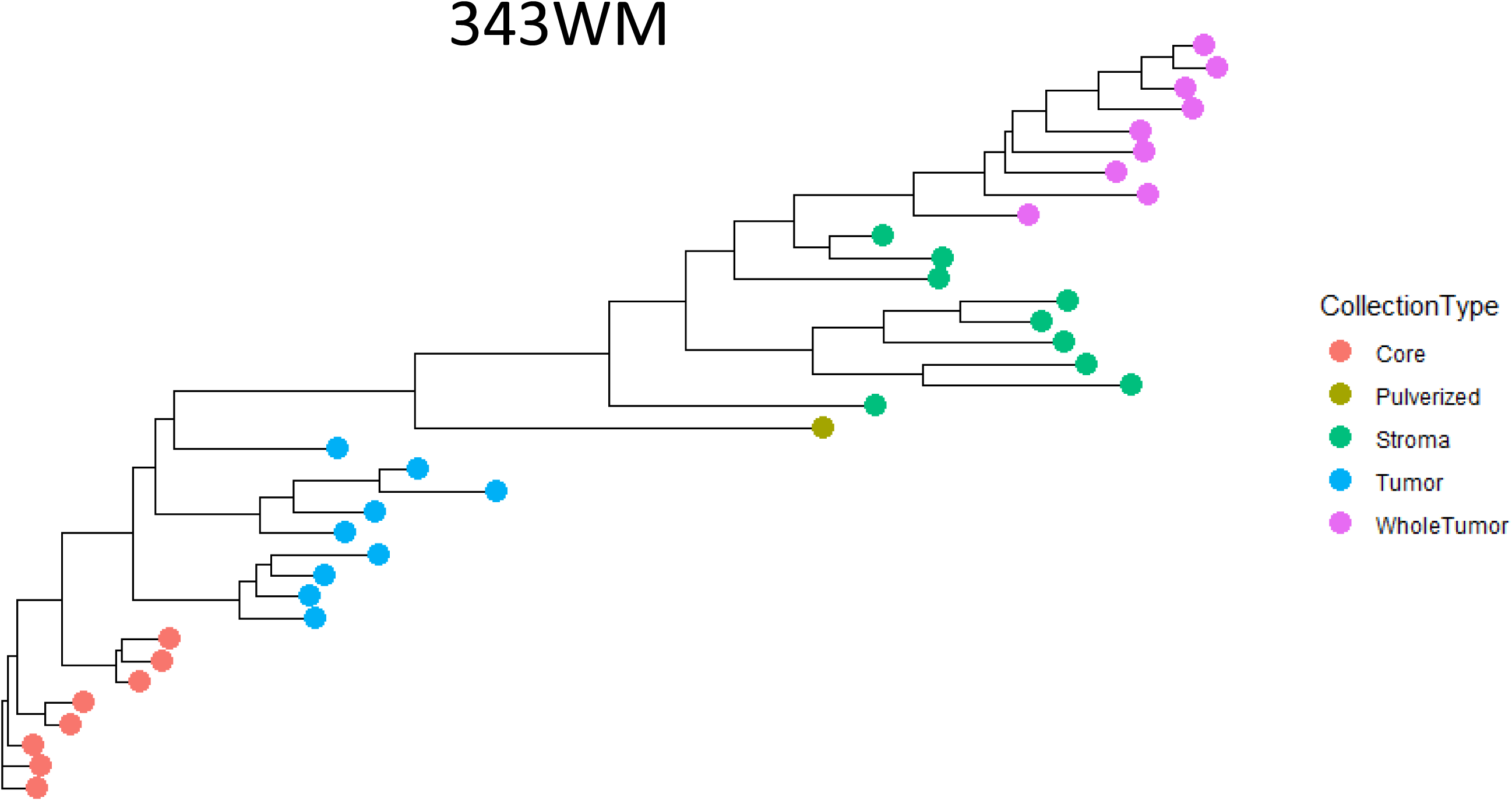
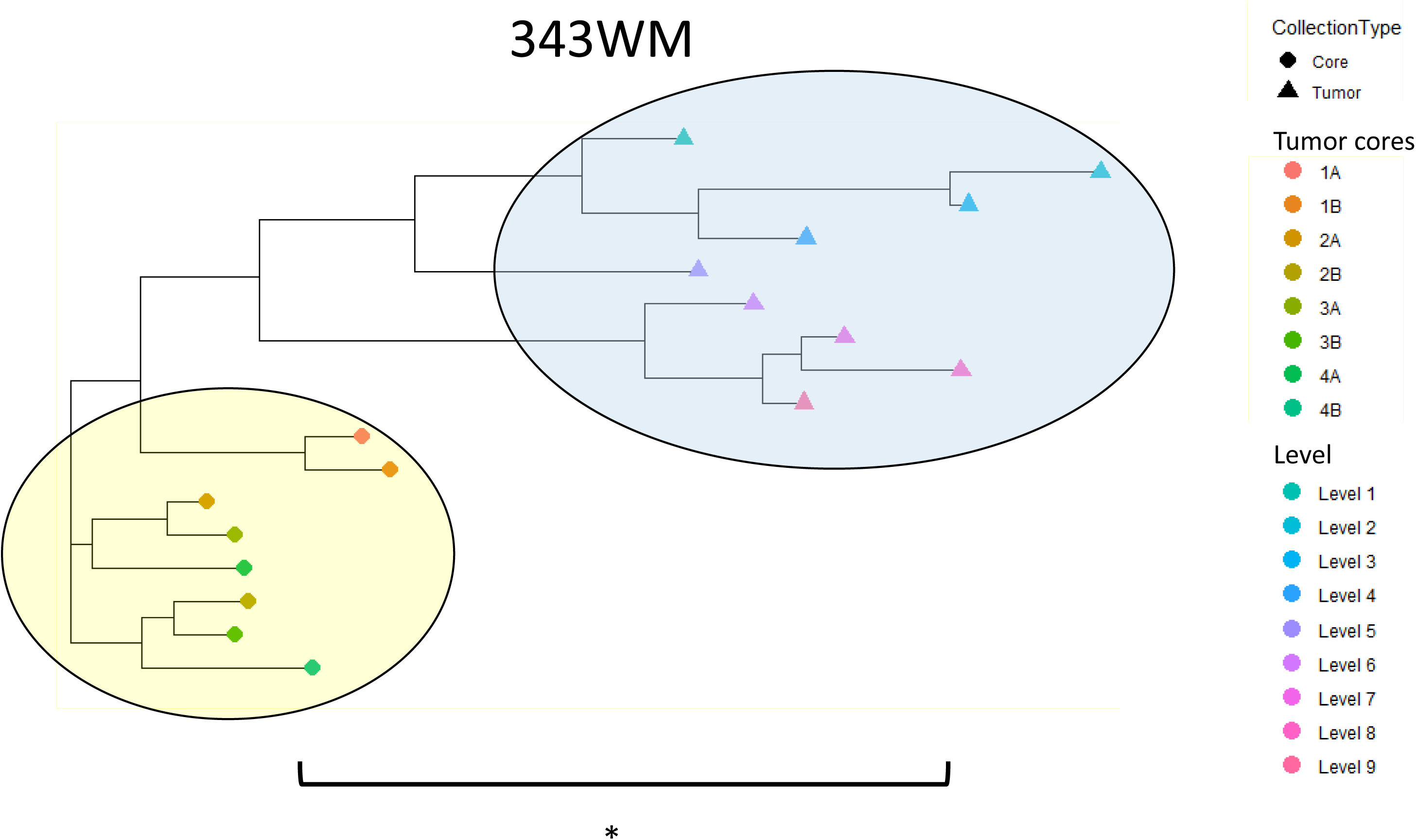
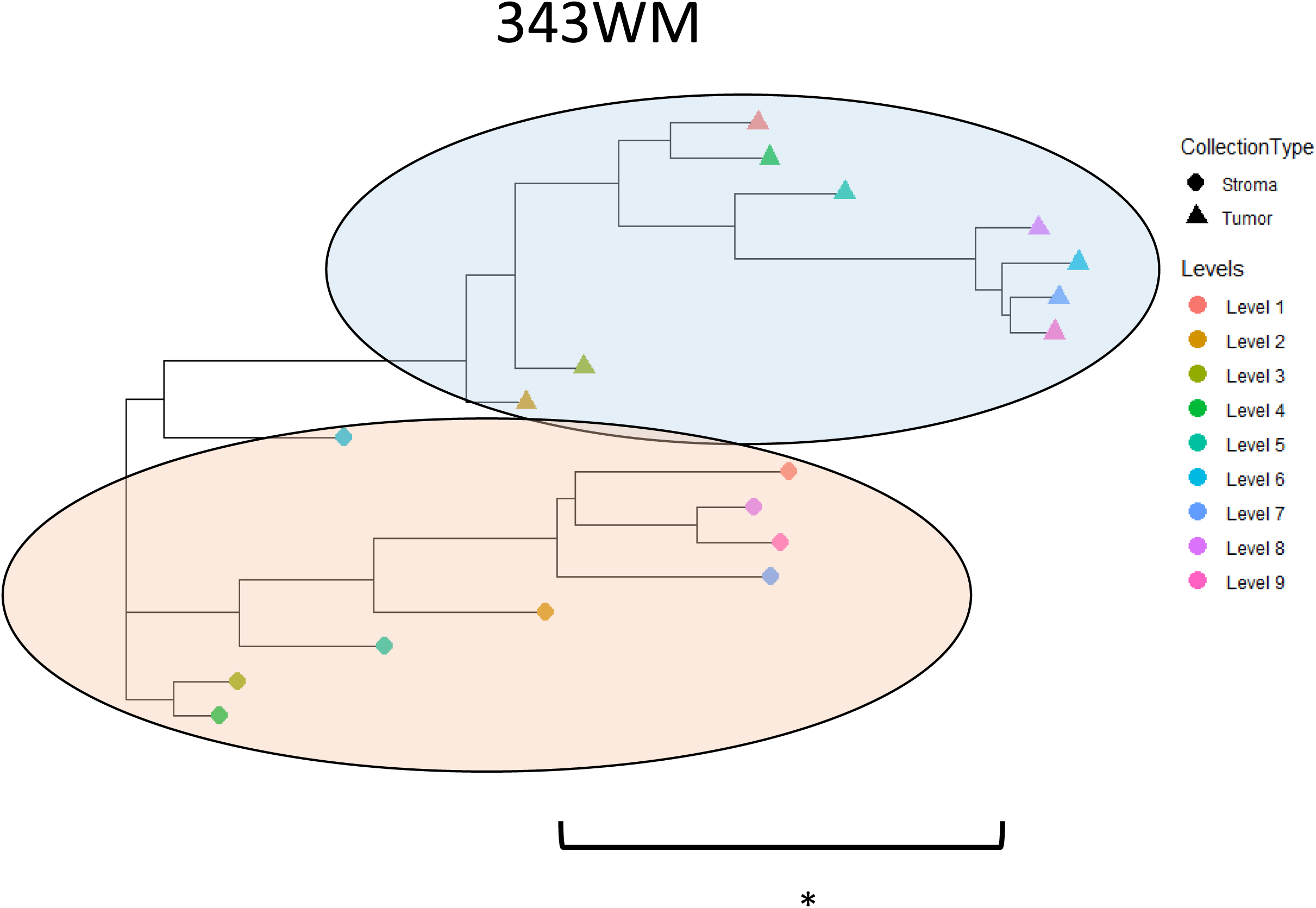
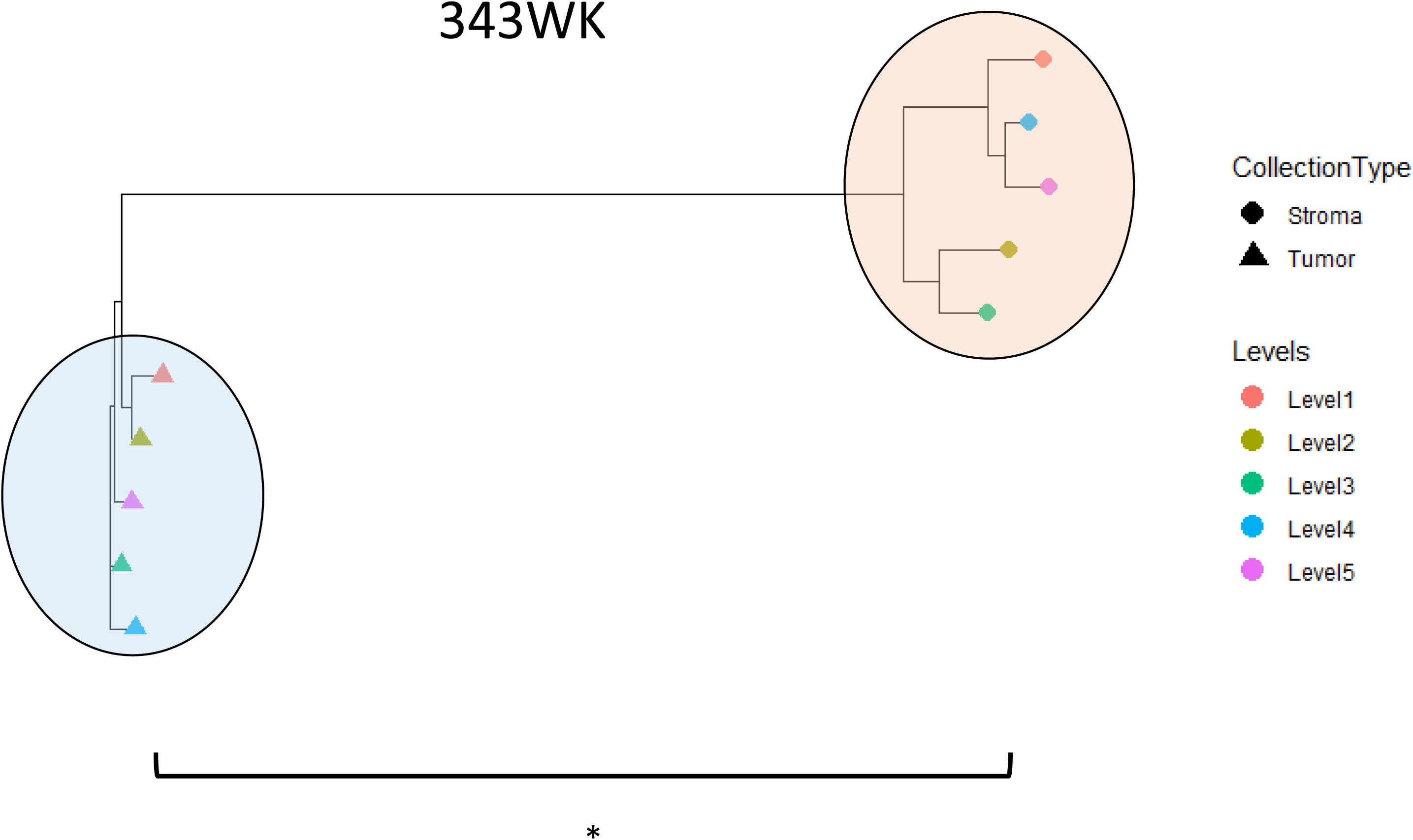
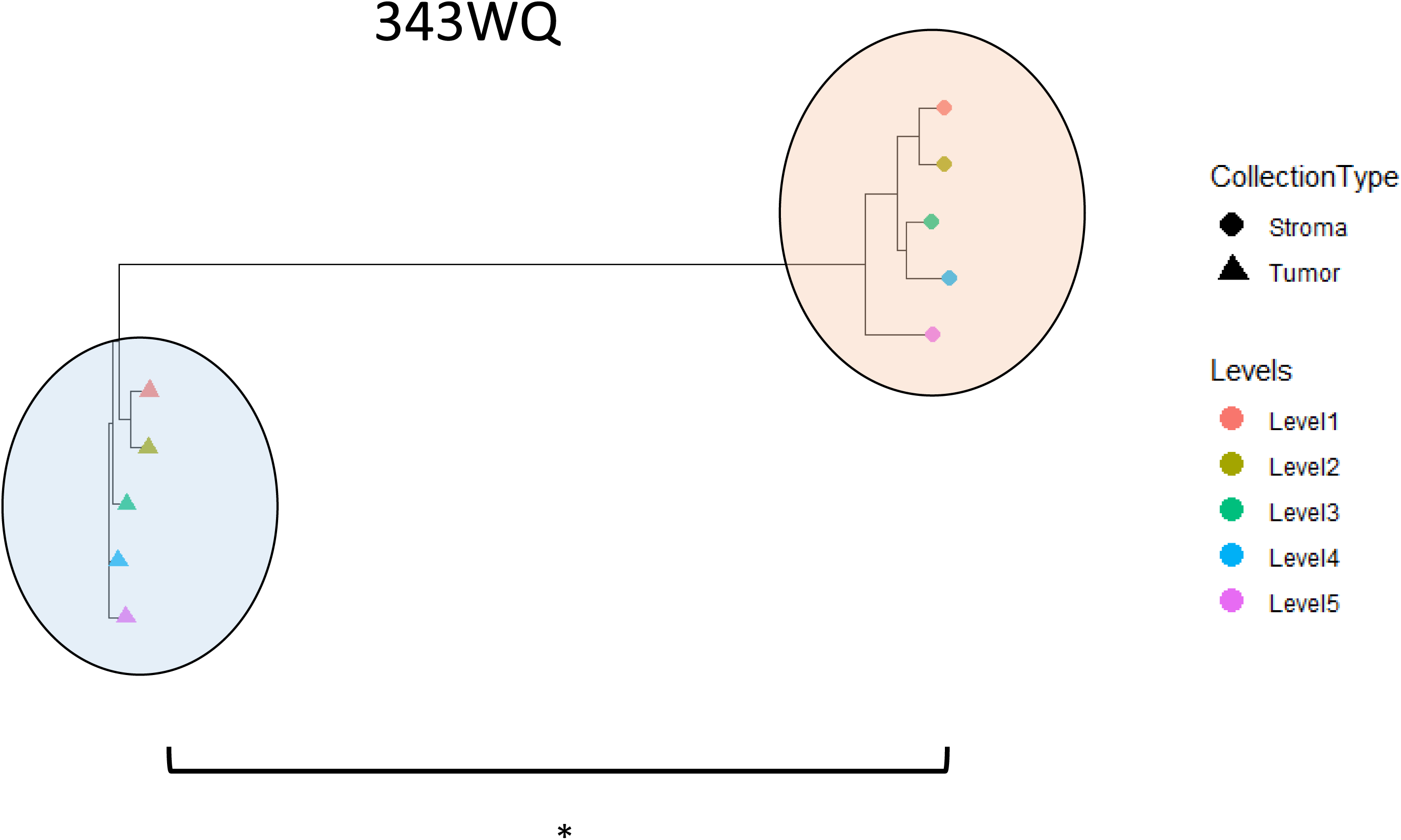
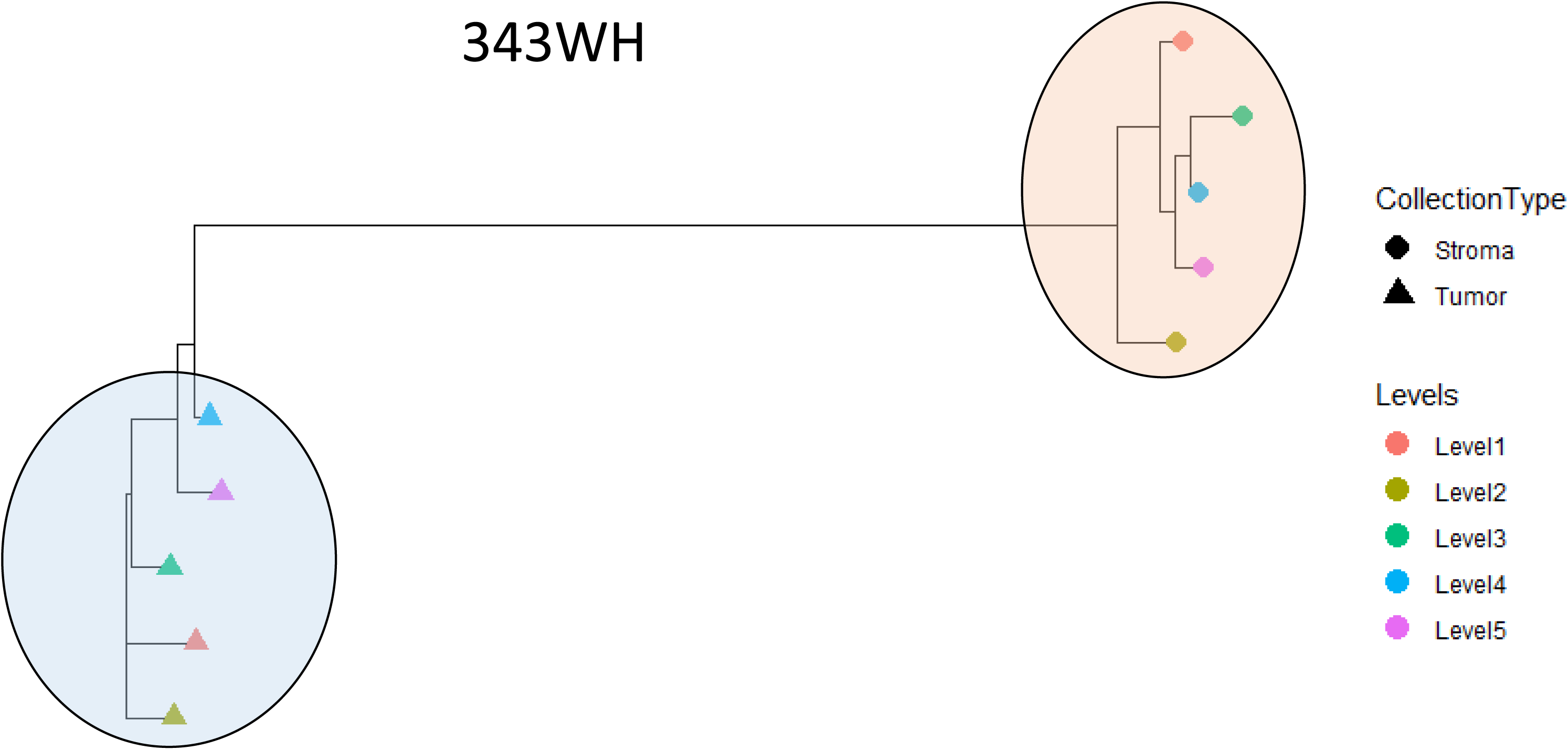
Patient-specific dendrograms depicting Spearman correlations between LMD enriched collections based on differentially expressed proteins. (A) Representative relatedness of all collection types from 343WM, including tumor cores, LMD enriched tumor epithelium, LMD enriched stroma, whole tumor, and cryopulverized tissue. (B) Relatedness of tumor cores versus LMD enriched tumor epithelium from 343WM. (C-F) Relatedness of LMD enriched tumor versus stroma from 343WM, 343WK, 343WQ, and 343WH, respectively. With the exception of the LMD enriched tumor cores vs LMD enriched tumor epithelium comparison in panel B which was calculated using proteins with a median absolute deviation (MAD) >0.5, all comparisons in panels A and C-F were made using proteins with MAD >1. Comparisons marked with (*) indicate a significant difference (p<0.01) between groups, with p-values of 1.264E-05, 0.0012, 0.0003372, 0.00097, and 0.2205 for the comparisons in panels B-F, respectively. The yellow, blue, and red ovals in panels B-F highlight the clusters of LMD enriched tumor cores, tumor epithelium, and stroma, respectively.

We next sought to explore molecular heterogeneity within each sample collection type using a phylogenic analysis approach depicting relatedness between samples, which was calculated based on similarities in protein abundance trends using Spearman correlations (Fig. 6; Supplemental Table 30). In this way, the correlations of samples within the same LMD collection type were calculated and then compared with those from another (as specified) LMD collection type. Comparison of enriched tumor and stroma collections in case 343WM (Fig. 6A) suggests that the proteome expression profiles of LMD enriched stroma are less heterogeneous from a phylogenic perspective than either of the LMD enriched tumor epithelium collections, and that the proteomic expression profile of whole tumor is more closely related to the LMD-enriched stroma. Focusing only the LMD enriched tumor epithelium collections (cores and cross sections) illustrates that (Fig. 6B) the degree of heterogeneity between all tumor cores is significantly less than the degree of heterogeneity observed between LMD enriched tumor collections (p<0.0001). Next, the correlations within and between the LMD enriched tumor epithelium and stroma samples were calculated for cases 343WM, 343WK, 343WQ, and 343WH. These four cases were selected as they each have equal numbers of LMD enriched tumor and stroma samples collected per case. In three of the four cases, the degree of heterogeneity between tumor epithelium samples was significantly less (p<0.01) than the heterogeneity observed in the stroma (Fig. 6C-F).

Striking examples of how regional tissue variations within a single tumor specimen can impact molecular subtype classification and measurement of proteomic markers of interest are highlighted for case 343WQ (Fig. 7), with similar findings seen across all cases (data not shown). The relative composition of tumor epithelium and stroma are generally unchanged throughout the depth of the specimen block (Fig. 7; left panel). The LMD enriched tumor and stroma collections across each of the 5 levels displayed consistent patterns of molecular abundances, correlations with HGSOC consensus molecular subtypes [4], and cellular admixture scores [13], though the magnitude of the correlations varied by level (Fig. 7; right panel). In contrast, the whole tumor collections displayed widely varied patterns of molecular abundances, subtype classifications, and cellular admixture scores across the 5 levels.

**Figure 7.**
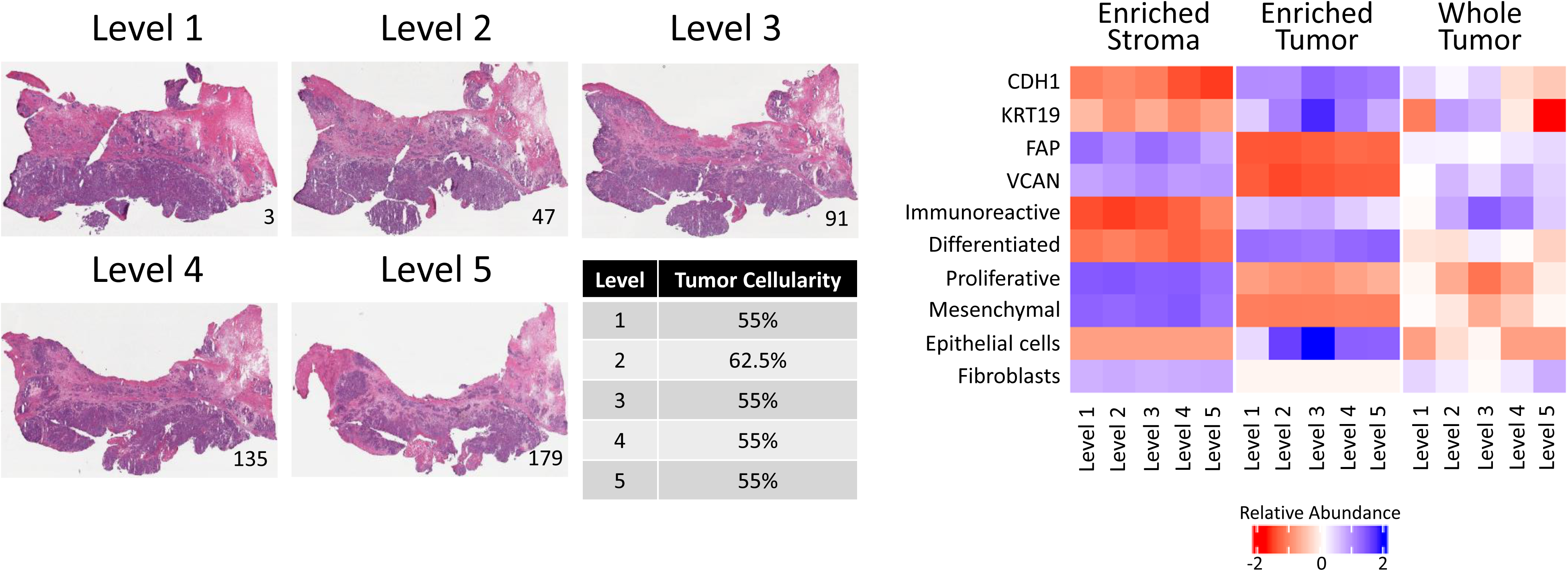
Representative case (343WQ) depicting variable molecular expression and subtype classification by level. H&E images show the tissue sections mounted on glass slides bounding the top of each level. The tissue section number is notated in the bottom right corner of each micrograph. The median tumor cellularity with relative standard deviation (%CV), molecular subtype, and protein abundances of representative tumor and stroma markers present in the whole tumor collections are indicated for each level. The median tumor cellularity calculated from review of multiple images per level with the %CV included in parenthesis (from Supplemental Table 1) are reported as percentages. Correlations with consensus molecular subtypes (Figure 5B and Supplemental Table 28) are shown. The Log_2_-transformed protein abundances are shown for tumor/epithelial markers (CDH1 and KRT19) and stroma markers (FAP and VCAN) (from Supplemental Table 7).

Finally, we selected three patients in our expanded cohort for whom complete cancer-related medical histories were available (Supplemental Figure 7) to analyze the heterogeneity of clinical biomarker expression between LMD levels within each patients’ specimen. Case 343WP was extremely sensitive to chemotherapy and surviving with no evidence of disease as of last follow-up, more than 5 years post-diagnosis. Cases 343WN and 343WQ had cancer-related deaths; these patients were moderately sensitive or refractory to chemotherapy, respectively. We generated a composite assessment of 70 genes or gene products analyzed in the clinical setting by Precision Therapeutics, Inc. (now Helomics (Pittsburgh, PA); a subsidiary of Predictive Oncology (Eagan, MN)), Caris Life Sciences (Irving, TX), Foundation Medicine (Cambridge, MA), Quest Diagnostics (Secaucus, NJ), and/or Ambry Genetics (Alisa Viejo, CA). Of these 70 markers, 31 were measured at the proteome level in our expanded cohort dataset. The variability of protein expression between levels for these markers was examined in cases 343WK, 343WQ, and 343WH, for whom the full complement of 5 levels/cases for all LMD collection types were available. Relative standard deviations (RSD) of protein expression were calculated and ranked to identify the top 5 most and least variable clinical markers (Supplemental Table 31). The most variable proteins between levels within a single patient’s specimen in our whole tumor samples (comparable with the type of tissue examined in the clinic) were ABL1, NF1, MSH6, MLH1, and PTEN. None of these were among the top 5 most variable in our LMD enriched tumor samples. TOP1, RAD50, EPCAM, SMARCA4, and APC were the least variable in whole tumor. TUBB3 and HRAS were among the top 5 most variable proteins in both LMD enriched tumor and LMD enriched stroma, while FBN1 was among the least variable in both LMD enriched compartments.

## Discussion

Ovarian cancers are typically diagnosed at an advanced stage after the accumulation of numerous molecular alterations [29]. In this study, we have demonstrated that LMD enrichment reveals extensive intratumoral proteogenomic heterogeneity, which has potential implications for diagnostic, prognostic and therapeutic predictive value, and is similar to what has recently been described in hepatocellular carcinoma [30].

In unsupervised hierarchical cluster analysis of proteins (Fig. 2A, 2C) and transcripts (Fig. 2B, 2D), all LMD enriched cell populations exhibited similar protein and transcript abundance trends. We found that many molecular alterations were negatively correlated with established diagnostic and prognostic signatures within discrete cell populations, such as confirmation of several mesenchymal marker genes as emerging from stroma cells in HGSOC tissues recently identified by Zhang *et al.* [31] (Fig. 5). Furthermore, we observe a profoundly unique transitioning of molecular subtype assignment based on sampling of the tumor microenvironment at the level of the proteome versus the transcriptome (Fig. 5A). These discrepancies are likely due to signature genes in molecular subtypes not being quantified in proteome-level data or exhibiting lower correlation with transcript-level abundance. We further note the correlation of protein and transcript abundance is higher in tumor epithelium versus stroma collections and is likely attributed to the secretory nature of stroma cells [32]. This is supported by the higher correlation of transcript abundance in stroma with protein abundance in whole tumor collections for case 343WM (Fig. 4), though this also likely represents overall tumor purity, as similar findings of higher correlation between stroma protein and whole tumor transcripts were not seen in 343WH (Supplemental Fig. 4). This was further underscored by our findings that the variance of proteins containing signal peptide sequences or classified as “extracellular” by Gene Ontology across all levels from each LMD enriched tumor and stroma was significantly greater (p<0.0001) than the variance of proteins lacking these classifications, and the lack of differential MUC16 protein abundance between these tissue compartments (Supplemental Fig. 3).

Comparison across proteomic platforms using proteins co-quantified by MS-based proteomics and RPPA revealed similar findings as seen above with the co-quantified proteins and transcripts in which there was general positive correlation within the same LMD enriched tumor epithelium or stroma tissues across all cases (Supplemental Fig. 5A, B). However, there was poor concordance between proteins measured in the whole tumor collections, often even within the same case (Supplemental Fig. 5C). While this is likely partially attributable to the limited number of features (54 proteins) used in this comparison [33], it largely recapitulates the findings that whole tumor collections reflect overall levels of tumor purity and the need for cell type enrichment for molecular characterization.

Our results suggest that molecular signatures developed to date have reflected variations in tumor purity that differentially impact outcome; specifically, lower tumor purity contributes to the assignment of tumors to poorer prognostic outcome (Fig. 5; Fig. 8). The predictive value for estimating patient outcome through molecular subtype assignment (using the molecular prognostic signatures identified by Konecny et al. [22]) showed a median survival difference of two years depending on whether the tumor epithelium or stroma and/or mixed tissue is sampled. Specifically, the mesenchymal subtype in the Konecny et al. study correlated with our LMD enriched stroma and had a median overall survival (OS) of approximately 24 months whereas the differentiated and proliferative subtypes correlated with our LMD enriched tumor and had a median OS of approximately 48 months [22]. Notably, the contribution of the stromal microenvironment to the mesenchymal signature has been recently described [31, 34, 35]. The observation that poor prognosis molecular subtypes (namely, mesenchymal subtype) are driven by the admixture of stroma cells supports the existence of a pathological ovarian stroma and support the proposed role of cancer-associated fibroblasts contributing to disease development and/or progression [36–39]. The presence of a large ovarian stromal compartment is an indicator of poor prognosis [40]. Tumors with more interceding stroma have lower resectability, decreasing the likelihood of achieving complete resection (R0) during surgical debulking leading to worse outcome [41, 42]. Gene signatures in the ovarian stroma have also been correlated with increased tumor invasiveness and metastatic spread to the blood and lymphatics [43]. However, assignment of the cryopulverized tissue in our study to a particular subtype was less clear due to profound signal averaging of the proteome (Fig. 5A). TCGA and other large-scale tumor characterization efforts to date have applied pre-determined tumor purity cutoff levels to include predominantly high purity tumors in their studies. As a result, molecular drivers of disease progression specific to prognostically poor/low purity tumors are frequently missed [4, 44], as recently realized for colon cancer [45] and gliomas [46].

We demonstrate the stromal distribution of many markers previously published as correlative with residual disease and/or suboptimal cytoreduction in ovarian cancer (Supplemental Fig. 6) [23, 24]. Among these, *FABP4* and *ADH1B* expression were associated with significantly higher levels of residual disease [23]. Expression and shedding of AKAP12 has been recently described in isogenic cell line models of paclitaxel resistance and elevation of AKAP12 transcript also correlated with decreased survival in HGSOC patients [47]. We observe AKAP12 transcript is elevated in stroma versus tumor epithelium (Supplemental Tables 5, 6, 7, 8) suggesting that elevation of AKAP12 in drug-resistant HGSOC cells may be due to chemotherapy-induced epithelial to mesenchymal transition and association with poor disease outcome may correlate with patients harboring lower-purity, likely mesenchymal-subtype tumors. *COL11A1* transcripts were also abundant in the stroma and some of the whole tumor collections (Supplemental Fig. 6), consistent with findings that COL11A1 is elevated in cancer-associated fibroblasts (CAFs) [37].

Extensive intratumor heterogeneity was observed in our study within each of the LMD enriched tumor and stroma compartments, as demonstrated by the tumor cores from case 343WM and between the spatially separated collection levels for each of the 10 HGSOC cases assessed. A comparison of the magnitude of heterogeneity within each LMD enriched tissue type revealed that overall there was a greater amount of heterogeneity in the stroma tissue relative to tumor epithelium (Fig. 6C-F). Though more related than the stroma collections, we demonstrate heterogeneity within the tumor epithelium resulting from locoregional differences, as replicate tumor cores displayed greater relatedness to each other than to LMD enriched tumor collections taken from the entire tissue area (Fig. 6A, B). Consistent with our enriched tumor and stroma cell findings, single cell RNA sequencing on HGSOC has previously revealed that individual cells of the same tissue type (tumor epithelium or stroma) differentially clustered with molecular subtypes due to unique gene expression profiles [48]. Further, heterogeneity within HGSOC molecular subtypes has been partially attributed to activation of distinct transcriptional factors serving as master regulators of the cellular phenotypes [49]. As mentioned, within the ovarian stromal microenvironment we observed the highest probability of classification as mesenchymal and/or immunoreactive subtypes (Fig. 5).

Compartmentalized expression of genes encoding known FDA-approved anticancer drug targets [21] was found in both the LMD enriched tumor epithelium and stroma collections (Supplemental Tables 23, 27). The proteins encoded by *VEGFA* and *FLT1* (also known as *VEGFR1*) are targets of bevacizumab, a monoclonal antibody often used as an anti-angiogenic therapy for treating HGSOC patients [50]. In our study, *VEGFA* transcript expression from case 343WM was significantly enriched in tumor epithelium whereas transcript expression of *FLT1*/*VEGFR1* was more abundant in LMD enriched stroma (Supplemental Table 23). Dysregulated VEGF/VEGFR expression has been demonstrated to contribute to epithelial ovarian cancer development and/or progression through increased vascularization and improved survival of endothelial cells via anti-apoptotic signaling in the newly formed vessels [51, 52], though bevacizumab targeting of VEGF/VEGFR has disputed levels of improvement on HGSOC patient survival [53]. PARP1 protein was consistently enriched in the tumor epithelium of 9 patients in our study (Supplemental Table 27). Olaparib, rucaparib, and niraparib are poly(ADP-ribose) polymerase (PARP) inhibitors used for treating HGSOC patients in either the recurrent or maintenance setting [54]. Conversely, MAP1A is a target of estramustine and PDGFRB is a target of sorafenib, regorafenib, and pazopanib; MAP1A and PDGFRB proteins were both found to be significantly enriched in LMD stroma.

Accurate diagnosis and estimation of prognostic outcome are of utmost interest to patients, so we analyzed the intratumor heterogeneity of protein expression for gene and/or protein markers that are routinely assessed in HGSOC patients. Specifically, we examined the variability of protein expression between each of the 5 levels per LMD collection type separately for cases 343WK, 343WQ, and 343WH and reported proteins that were most and least variable within each LMD collection type (Supplemental Table 31). Our whole tumor samples represent the types of samples analyzed in the clinical setting. ABL1 was among the top 5 most variable proteins in whole tumor samples within each patient’s tumor. ABL1 is a target of the drug Regorafenib which is currently FDA-approved for treatment of colorectal and stomach cancers [55] and has been the subject of recent clinical trials for treatment of recurrent epithelial ovarian cancers. Interim clinical trial results reported Regorafenib to be inefficacious in improving progression free survival, overall survival, or overall response rate. Our results suggest that the clinical inefficacy of Regorafenib for treating ovarian cancers could, at least in part, be attributed to the high intratumoral heterogeneity of expression of ABL1 or other related proteins. MSH6 and MLH1 were also among the top 5 most variable proteins in this analysis. MSH6 and MLH1 are biomarkers of mismatch repair and microsatellite instability and are used as prognostic indicators of improved survival and clinical response to platinum-based chemotherapy [56]. The high levels of intratumoral heterogeneity of MSH6 and MLH1 in our study suggest that the predictive efficacy of platinum-based chemotherapies may not adequately represent true patient response due to the potential for sampling variations of regional protein expression within a tumor biopsy. TUBB3 is the target of the drug Ixabepilone, FDA-approved for treatment of breast cancer [55] and the subject of recent clinical trials for potential use in patients with recurrent or persistent platinum- and taxane-resistant ovarian cancer [57], which showed promising results. Surprisingly, TUBB3 was among the top 5 most variable proteins in both LMD enriched tumor and LMD enriched stroma in our analysis.

In conclusion, we demonstrate a critical need to account for cellular subtype and regional TME proteogenomic heterogeneity in cancer molecular profiling efforts that will substantially enable in-depth characterization of spatially distributed subclonal cell populations that have underappreciated roles in driving carcinogenesis. Further work to deconstruct the TME will aid clinical diagnosis, improve the efficacy of therapeutic intervention, and increase capabilities to identify druggable molecular markers of disease development and progression.

## Materials and Methods

### Tissue specimens

Surgically resected fresh-frozen tumor tissue specimens were obtained from ten patients with HGSOC of primary ovarian or tubo-ovarian origin. These comprised a mixture of chemotherapy-naïve specimens and specimens obtained following neoadjuvant chemotherapy (NACT) from patients with advanced stage (stage III or IV) disease. The specific tissues used for the study were from ovary, omentum, and pelvic or adnexa masses. The tissue blocks were sectioned by cryotome into 170 - 210 consecutive 10 µm thin tissue sections (1.7 - 2.1 mm total depth) which were placed on PEN membrane slides (Leica Microsystems). Representative sections were mounted on charged glass slides and stained with hematoxylin and eosin (H&E) after every 10 PEN membrane slide sections (100 µm), except for case 343WM where representative H&E slides were generated after every 20 PEN membrane slide sections (200 µm).

### Laser microdissection

The specimen obtained from case 343WM was uniquely prepared for laser microdissection (LMD) and proteogenomic analysis of spatially separated tumor “cores” which represent defined sub-populations of tumor epithelium that were collected as technical replicate areas for comparison with tumor epithelium collected from larger whole slide areas. Four regionally separated areas within the tissue were laser microdissected to enrich tumor epithelium (LMD7, Leica Microsystems) from each of 100 slides for proteomics or 50 slides for transcriptomics. The cores were pooled from a cross-sectional area of approximately 1 mm^2^ collected by LMD per slide for a depth spanning the entire block (1 mm x 1 mm x 2 mm). An adjacent 1 mm^2^ was microdissected to serve as a replicate from each core; in total each core plus their respective adjacent replicate regions covered an approximate 2 mm^3^ aggregate volume of tumor epithelium. The distance between the center of each tumor core and the center of the nearest neighboring core was approximately 4 mm. For a set of 9 sections (200 µm apart) each for proteomics and transcriptomics, all remaining tumor epithelium (between 20 - 44 mm^2^/section) and stroma (44 - 80 mm^2^/section) were collected by LMD after the core regions were collected. Whole tumor tissue was collected from a second set of 9 sections (96 - 155 mm^2^/section), where only regions containing necrosis, blood, and fat were excluded.

For the remaining 9 HGSOC cases, the slides from each block were separated into five equal levels (quintiles) assigned by depth within the block. Slides designated for MS proteomics (n=9 patients, 5 levels each), reverse phase protein array (RPPA; n=8 patients, 5 levels each, plus 8 levels from 343WM), and transcriptomics (n=2 patients, 1 level each) within each level were interlaced as much as possible.

All slides were H&E stained prior to LMD. For the slides microdissected for mass spectrometry (MS) proteomics, phosphatase inhibitors (Sigma Aldrich) were added to the 70% ethanol fixative and the first DEPC water; slides for transcriptomics were stained with RNase inhibitors (ProtectRNA; Sigma Aldrich) added into all aqueous solutions. Slide preparation for RPPA did not involve staining with eosin after the hematoxylin but included color development in Scott’s Tap Water (Thermo Fisher Scientific) and two final drying steps in xylenes. Protease inhibitors (Roche) were added to all RPPA staining solutions except for the 100% ethanol and xylene washes.

Within each quintile level per specimen block LMD was used to enrich cross-sectional areas of 40 mm^2^ and 15 mm^2^ (LMD enriched tumor epithelium, LMD enriched stroma, and whole tumor) for analysis via liquid chromatography-tandem mass spectrometry (LC-MS/MS) and reverse phase protein microarray (RPPA), respectively. For two cases (343WH and 343WB), a cross-sectional area of 25 mm^2^ of each LMD enriched tumor, LMD enriched stroma, and whole tumoe was collected from one level for RNA sequencing (RNA-seq). Microdissected tissue was collected into LC-grade water (Fisher Scientific), a tissue extraction/lysis buffer, or Buffer RLT (Qiagen) for analysis via LC-MS/MS, RPPA, and RNA-seq, respectively. The RPPA extraction/lysis buffer consisted of a 1:1 mixture of Tissue Protein Extraction Reagent (T-PER; Pierce), Novex 2x Tris-Glycine SDS Sample Buffer (Invitrogen), plus 2.5% *v*/*v* 2-mercaptoethanol which was added to tissue at a ratio of 3 µl buffer per 1 mm^2^ LMD tissue. For RPPA, each slide spent no more than 30 min dwell time on the LMD microscope. Additional slides were collected into new tubes. Collected tissue was briefly centrifuged and frozen at -80 °C until analysis.

### OracleBio image analysis

Pre- and post-LMD images were collected using the Aperio ScanScope XT slide scanner (Leica Microsystems). Image analysis was performed by OracleBio using the Indica Labs HALO platform. Post-LMD images from 343WM were used to develop classification algorithms for identification of the “dissection area” and “all remaining tissue”. A separate algorithm was developed which involved co-registration of post-LMD images with corresponding adjacent H&E-stained sections on glass slides for detection and quantification of the cell nuclei abundance within the LMD regions. The size and number of cells and nuclei collected in each LMD collection was determined using matched sets of reference glass H&E sections and PEN membrane slides following LMD enrichment of tumor (n=15) and stroma (n=6) cell populations.

### Peptide preparation for TMT LC-MS/MS

LMD tissue in LC-grade water was dried, re-suspended in 100 mM triethylammonium bicarbonate (TEAB)/10% acetonitrile (ACN), and digested using SMART trypsin (1 µg/30 mm^2^ tissue; Thermo Fisher) and pressure cycling technology (PCT, Pressure Biosciences, Inc.) [58]. For 343WM, the remaining tissue in the OCT block was washed with water and cryopulverized in liquid nitrogen in a mortar and pestle. The cryopulverized tissue was re-suspended in 100 mM TEAB/10% ACN and digested by PCT using SMART trypsin (1 µg protease/ 30 mm^2^ tissue). Peptide concentrations from trypsin digests were determined using the bicinchoninic acid assay (Pierce BCA). Tryptic peptides were labeled with isobaric Tandem Mass Tags (TMT) according to the manufacturer’s instructions using the TMT 10-Plex Kit or TMTpro 16-Plex Kit from Thermo Fisher (Supplemental Table 4). The TMT 10-Plex reagents and TMTpro 16-plex reagents were used for labeling samples from case 343WM the remaining nine cases, respectively, to generate patient-specific TMT plexes. Briefly, peptide samples were individually mixed with respective TMT reagents for 1 h at room temperature, then quenched using 5% hydroxylamine. Empty TMT channels were filled using equal amounts of peptide mixed from each of the 5 levels of the whole tumor collections in the respective patient-specific plex.

Each TMT-10 or TMTpro-16 multiplex set of samples were loaded onto a C-18 trap column in 10 mM NH_4_HCO_3_ (pH 8.0) and fractionated by basic reversed-phase liquid chromatography (bRPLC) into 96 fractions through development of a linear gradient of acetonitrile (0.69% acetonitrile/min). For 343WM, 36 concentrated fractions were generated by pooling in a serpentine manner, each of which were analyzed for LC-MS/MS. For the remaining 9 cases, 24 concentrated fractions were generated following bRPLC which were individually analyzed by LC-MS/MS.

### Liquid chromatography-tandem mass spectrometry

The TMT-10 or TMTpro-16 sample multiplex bRPLC fractions were analyzed by LC-MS/MS employing a nanoflow LC system (EASY-nLC 1200, ThermoFisher Scientific, Inc.) coupled online with an Orbitrap Fusion Lumos Tribrid MS (ThermoFisher Scientific, Inc.). In brief, each sample was loaded on a nanoflow HPLC system outfitted with a reversed-phase trap column (Acclaim PepMap100 C18, 2 cm, nanoViper, ThermoFisher Scientific, Inc) and a heated (50 °C) reversed-phase analytical column (Acclaim PepMap RSLC C18, 2 μm, 100 Å, 75 μm × 500 mm, nanoViper, ThermoFisher Scientific, Inc) connected online with an Orbitrap mass spectrometer. Peptides were eluted by developing a linear gradient of 2% mobile phase B (95% acetonitrile with 0.1% formic acid) to 32% mobile phase B within 120 min at a constant flow rate of 250 nL/min. High resolution (R=60,000 at *m/z* 200) broadband (*m/z* 400 – 1,600) mass spectra (MS) were acquired from which the top 12 most intense molecular ions in each MS scan were selected for high-energy collisional dissociation (HCD, normalized collision energy of 38%) acquisition in the orbitrap at high resolution (R=50,000 at *m/z* 200). Monoisotopic precursor selection mode was set to “Peptide” and MS1 peptide molecular ions selected for HCD were restricted to z = +2, +3 and +4. The RF lens was set to 30% and both MS1 and MS2 spectra were collected in profile mode. Dynamic exclusion (t=20s at a mass tolerance=10 ppm) was enabled to minimize redundant selection of peptide molecular ions for HCD.

### Quantitative proteomic data processing pipeline for global proteome analysis

Peptide identifications were generated by searching the .raw data files with a publicly-available, non-redundant human proteome database (Swiss-Prot, Homo sapiens, Proteome UP000005640, 20,257 sequences, downloaded 12-01-2017; http://www.uniprot.org/) appended with porcine trypsin (Uniprot: P00761) sequences using Mascot (Matrix Science) and Proteome Discoverer (Thermo Fisher Scientific). Identification, normalization, and Log_2_ transformation of PSMs was performed as previously described [58] for calculation of protein-level abundances.

### RNA sequencing

Tissue collected by LMD in Buffer RLT was purified using the RNeasy Micro Kit (Qiagen) per the manufacturer’s instructions. RNA concentrations were determined by fluorescence (Qubit HS and BR kits, Thermo Fisher). RNA integrity numbers (RIN) were calculated using the RNA 6000 Pico Kit 2100 Bioanalyzer (Agilent). All RNA was high-quality with RIN values of 7.2 or greater, except for two samples which had RIN values of 4.9 and 6.5.

RNA samples were reverse transcribed from 10 ng input using the SuperScript VILO cDNA Synthesis Kit. Barcoded cDNA libraries containing 5-6 LMD samples plus a Universal Human Reference RNA (UHR) standard (Stratagene) were prepared on the Ion Chef System using the Ion Ampliseq Chef DL8 materials and the Ion AmpliSeq Transcriptome Human Gene Expression Panel Chef-ready Kit (Thermo Fisher). Libraries were then purified via solid phase reversible immobilization (SPRI) using AMPure XP beads (Beckman Coulter) to remove fragments less than 100 bp, quantified by qPCR (TaqMan Quantitation Kit; Applied Biosystems), and 25 µl of 100 pM diluted library was used for templating, amplification via emulsion PCR, and loaded onto Ion 550 chips on the Ion Chef System.

Sequencing was performed on the Ion Torrent S5 XL (Thermo Fisher) and mapped to the hg19 human reference transcriptome (hg19_Ampliseq_Transcriptome_21K_v1). Successful sequencing runs achieved ≥18M reads/sample (with one exception) and 169-234X mean coverage depth. Per the Torrent Suite Software (Torrent Suite v5.8.0), the number of reads aligning to a given gene target represents an expression value referred to as “counts”. The read count per million mapped reads (RPM) for each barcoded sample was calculated by the software as read count x 10^6^ / total number of mapped reads. Normalized RPM-level transcript abundances were calculated relative to the average RPM abundance quantified across all samples for a given transcript and Log_2_ transformed.

### Bioinformatic and statistical analyses

Unsupervised analyses were performed using proteins or transcripts exhibiting a median absolute deviation (MAD) >0.5 across all samples and clustered by Pearson correlation as heatmaps using Clustvis (version 1.2.0) in R (version 3.6.2). Differential analyses of global proteome and transcript data matrixes was performed using LIMMA (version 3.8, [59]) in R (version 3.6.2); features satisfying a LIMMA p-value <0.01 and exhibiting a Log_2_ fold-change cut-off ±1 were prioritized for downstream analyses. Cell type enrichment analyses were performed using RPM-level RNA-seq data in xCell (http://xcell.ucsf.edu/; version 1.1.0 [60]). xCell cell type signature scores of interest were categorized by relative rank from highest to lowest spanning a range of 1 to -1 to enable co-visualization with similarly categorized transcript and protein abundance for candidates of interest. Boxplots of relative protein abundances or cell type signature scores were generated using Ggplot2 (version 3.2.1). Random forest probabilities for classification of consensus molecular subtypes using either global proteome or transcriptome data were calculated using consensusOV (version 1.8.1). Dendrograms were generated using Ggtree (version 2.0.1) using features with MAD>1 across individual patient samples, with the exception of the comparison between tumor cores and LMD enriched tumor from case 343WM, which used features with MAD>0.5. Tree structure was determined by neighbor-joining tree estimation using Ape (version 5.3) and Spearman correlations between samples were calculated. Molecular subtype classifications were compared by Spearman correlation using MedCalc (version 19.0.7). Functional pathway inference and drug targets were assessed using Ingenuity Pathway Analysis. The heatmap in Fig. 7 was generated using ComplexHeatmap (version 2.2.0); the heatmaps in Supplementary Fig. 4 and 5 were generated using Morpheus (https://software.broadinstitute.org/morpheus/; version 1.0-1).

### Reverse phase protein microarray

Using an Aushon 2470 arrayer (Aushon BioSystems, Billerica, MA) equipped with 185 μm pins, cell lysates along with internal reference standards were immobilized onto nitrocellulose-coated slides (Grace Bio-labs, Bend, OR) in technical replicates (n=3). To assess the amount of protein in each sample and for normalization purposes, selected arrays were stained with Sypro Ruby Protein Blot Stain (Molecular Probes, Eugene, OR) following manufacturer’s instructions. Immunostaining was performed as previously described [61, 62]. In brief, prior to antibody staining, arrays were first treated with Reblot Antibody Stripping solution (Chemicon, Temecula, CA) for 15 minutes at room temperature, washed with PBS and incubated I-block (Tropix, Bedford, MA). To reduce unspecific binding between endogenous proteins and the detection system, arrays were then probed with 3% hydrogen peroxide, an avidin/biotin blocking system (Dako Cytomation, Carpinteria, CA), and an additional serum free protein block (Dako Cytomation, Carpinteria, CA) using an automated system (Dako Cytomation, Carpinteria, CA). Each array was then probed with one polyclonal or monoclonal primary antibody targeting a protein of interest. Primary antibodies’ specificity was previously assessed on commercial cell lysates or human tissue lysates by western blotting [63]. The criteria for validation were the presence of a single band of correct molecular weight in the positive control sample and the absence of a band in the negative control. Antibodies targeting phosphorylated epitopes were additionally validated by ligand induction. A subset of antibodies was further validated against peptide competition. Arrays were probed with total of 281 antibodies targeting native and/or post-translationally modified proteins. Primary antibodies were recognized by a biotinylated anti-rabbit (Vector Laboratories, Inc.) or anti-mouse secondary antibody (Dako Cytomation, Carpinteria, CA). Signal amplification was achieved using a commercially available tyramide-based avidin/biotin amplification system (Dako Cytomation, Carpinteria, CA) coupled with fluorescent detection using the streptavidin-conjugated IRDye680 dye (LI-COR Biosciences, Lincoln, NE), according to the manufacturer’s instructions. Selected arrays were incubated with the secondary antibodies alone to capture background and unspecific signal. Images were acquired using a laser scanner (TECAN PowerScanner, Mönnedorf, Switzerland) and analyzed using a commercially available software (MicroVigene V5.1.0.0; Vigenetech, Carlisle, MA). In brief, the software performed spot finding and subtraction of the local background/unspecific signal. Finally, each sample was normalized to the corresponding amount of protein derived from the Sypro Ruby stained slides and triplicates were averaged.

## Acknowledgements

This study was supported in part by the U.S. Department of Defense - Uniformed Services University of the Health Sciences (HU0001-16-2-0006 and HU0001-16-2-0014) and the Ovarian Cancer Research Program from the Congressionally Directed Medical Research Program (W81XWH-16-2-0038). The authors would like to acknowledge Lorcan Sherry and Mark Anderson from OracleBio Limited (Scotland, UK) for histopathology image analysis support. The authors would like to acknowledge Victoria Olowu, Marshé Edwards, Fred Park, Persus Akowuah, Salma Eltahir, Katherine Zhou, Domenic Tommarello, Pang-Ning Teng, Katlin Wilson, Sakiyah TaQee, and Tamara Abulez for histopathology and proteomic sample preparation and informatic analysis support.

## Author Contributions

Contributed to conception: NWB, TPC. Contributed to experimental design: ALH, NWB, TPC. Contributed to identification and acquisition of the patient specimens: GG, DM, JO, GLM, TPC. Contributed to acquisition, analysis and/or interpretation of data: ALH, NWB, WB, SMM, BLH, KAC, MZ, VC, JL, TL, JO, DM, GG, CR, BB, UNMR, EFP, GLM, TPC. Drafted and/or revised the article: ALH, NWB, YC, KO, AKS, KMD, CDS, UNMR, MP, EFP, GLM, TPC. Acquired funding for the research: YC, GLM, TPC. All authors read and approved the final manuscript.

## Disclaimer

The views expressed herein are those of the authors and do not reflect the official policy of the Department of Army/Navy/Air Force, Department of Defense, or U.S. Government.

## Supplemental Table Legends

**Supplemental Table 1. Clinical features and manual pathology assessment of tumor purity throughout the depth of the HGSOC patient specimen block.** Representative H&E stained glass slides from were examined at ∼200 µm intervals for case 343WM or ∼100 µm intervals for the additional 9 cases by a board-certified pathologist for percent by area estimation of tumor cellularity, necrosis, stroma, normal ovarian epithelium, lymphocytes, and polymorphonuclear leukocytes (PMN). Multiple images per level per case were reviewed and the median tumor cellularity and median stroma cellularity are reported with corresponding percent coefficient of variation (%CV) reported in parentheses. Abbreviations: NACT= neoadjuvant chemotherapy; FT= fallopian tube; Ov= ovary; ca= cancer; NOS= not otherwise specified.

**Supplemental Table 2. Depiction of full study cohort.** Numerical values indicate the number of LMD tissue sections that were used for each collection.

**Supplemental Table 3. LMD enriched tumor epithelium and stroma cell areas acquired by laser microdissection and automated analyses of collected regions of interest (ROI).** ROI are denoted as millimeter squared (mm^2^) areas, cell nuclei as µm^2^ areas or as total cell counts derived from cellular nuclei measured in LMD enriched tumor or stroma ROIs.

**Supplemental Table 4. TMT Plexing Strategy.** For 343WM, tryptic peptides were labeled with isobaric Tandem Mass Tags (TMT) according to the manufacturer’s instructions using the TMT 10-Plex Kit from Thermo Fisher for a total of 4 plexes. 30 µg peptides from the 8 cores, 9 non-discrete whole tumor collections, and cryopulverized tissue sample were individually mixed with respective TMT reagents for 1 h at room temperature, then quenched using 5% hydroxylamine. 10 µg peptides/sample were similarly TMT labeled from the 9 discrete collections each of remaining LMD-enriched tumor epithelium or stroma. For the remaining 9 cases, digested peptide samples were labeled in patient-specific plexes using the TMTpro 16-Plex Kit (Thermo Fisher). Specifically, 5 µg peptide from up to 5 levels/patient each of LMD enriched tumor epithelium, LMD enriched stroma, and whole tumor collections were incubated with the TMTpro reagents per plex for a total of 80 µg/plex and 1 plex/patient. To replace samples from which <5 µg peptide was recovered, empty TMT channels were filled using equal amounts of peptide mixed from each of the 5 levels of the whole tumor collections.

**Supplemental Table 5. Global protein matrix.** Log_2_ transformed fold-change abundances of 6,053 proteins imputed across all samples measured in case 343WM.

**Supplemental Table 6. Global transcriptome matrix.** Normalized abundances of 20,535 RNA transcripts measured in case 343WM calculated relative to the average RPM abundance quantified across all samples for a given transcript, as reported by the Torrent Suite (v5.8.0) software, before Log_2_ fold-change transformation.

**Supplemental Table 7. Global protein matrix for expanded cohort (n=9 patients).** Log_2_ transformed fold-change abundances of 6,199 proteins imputed across all samples (n=123).

**Supplemental Table 8. Global transcriptome matrix for expanded cohort (n=2 patients).** Normalized abundances of 19,758 RNA transcripts calculated relative to the average RPM abundance quantified across all samples (n=6) for a given transcript, as reported by the Torrent Suite (v5.8.0) software, before Log_2_ fold-change transformation.

**Supplemental Table 9. Unsupervised analysis of protein abundance (median absolute deviation > 0.5) measured in case 343WM.**

**Supplemental Table 10. Unsupervised analysis of transcript abundance (Log_2_ fold-change abundances with median absolute deviation > 0.5) measured in case 343WM.**

**Supplemental Table 11. Unsupervised analysis of protein abundance (median absolute deviation > 1) measured in the expanded cohort (n=9 cases).**

**Supplemental Table 12. Unsupervised analysis of transcript abundance (median absolute deviation > 1) measured in the expanded cohort (n=2 cases).**

**Supplemental Table 13. Cell type enrichment analyses as performed using RPM-level RNA-Seq data for case 343WM and default settings in xCell (**http://xcell.ucsf.edu/**, [60]).**

**Supplemental Table 14. Cell type enrichment analyses as performed using protein-level TMT LC-MS/MS data from the expanded cohort (n=9 cases) and default settings in xCell (**http://xcell.ucsf.edu/**, [60]).**

**Supplemental Table 15. Median absolute deviation (MAD) of LMD-enriched samples expressing or lacking signal peptide sequences and extracellular classification (n=4 cases).** P-values indicate the reliability of the presence or absence of a signal peptide or extracellular classification within the indicated LMD-enriched tissue across all levels/case. 9 levels were used for comparison for case 343WM, while 5 levels/case were used each for cases 343WK, 343WQ, and 343WH.

**Supplemental Table 16. Co-quantified proteins and transcripts from case 343WM.** Log_2_ transformed fold-change abundances of 5,742 imputed proteins that were co-measured at the transcriptome level from case 343WM.

**Supplemental Table 17. Co-quantified proteins and transcripts in the expanded cohort (n=2 patients).** Log_2_ transformed fold-change abundances of 5,379 imputed proteins that were co-measured at the transcriptome level.

**Supplemental Table 18. Information for all antibodies used for RPPA analysis.**

**Supplemental Table 19. RPPA abundance values for protein and phosphoprotein targets in LMD enriched tumor, LMD enriched stroma, and whole tumor samples from n=9 HGSOC cases.**

**Supplemental Table 20. Spearman correlations for proteins measured by TMT LC-MS/MS and RPPA.**

**Supplemental Table 21. Top altered canonical pathways in laser microdissection-enriched tumor epithelium versus stroma identified using Ingenuity Pathway Analysis.** Comparison of the most differential pathways and diseases/functions between LMD enriched tumor and stroma was determined using pairwise supervised analysis of proteomic and transcriptomic data with LIMMA adjusted p-value < 0.01 exhibiting a Log_2_ fold-change cut-off ± 1.

**Supplemental Table 22. Top altered diseases and functions in laser microdissection-enriched tumor epithelium versus stroma identified using Ingenuity Pathway Analysis.** Comparison of the most differential diseases/functions between LMD enriched tumor and stroma was determined using pairwise supervised analysis of proteomic and transcriptomic data with LIMMA adjusted p-value < 0.01 exhibiting a Log_2_ fold-change cut-off ± 1.

**Supplemental Table 23. Significantly differentially expressed transcript alterations between LMD enriched tumor epithelium versus stroma from case 343WM identified using Ingenuity Pathway Analysis (IPA) of known gene targets of FDA-approved anticancer drugs.** Comparison of prevalent drug targets present in LMD enriched tumor versus stroma was determined using pairwise supervised analysis of proteomic and transcriptomic data with LIMMA adjusted p-value < 0.01 exhibiting a Log_2_ fold-change cut-off ± 1. The complete list of all differentially expressed genes that are known drug targets from our proteomic-level and transcriptomic-level data was compared with a list of 150 FDA-approved anticancer drugs as of 2014 analyzed in a study by Sun et al [21].

**Supplemental Table 24. Proteomic-level data for 381 proteins which displayed the same pattern of enrichment across all 9 cases and which had Log_2_ fold-changes > ±1 used as input for Ingenuity Pathway Analysis.**

**Supplemental Table 25. Top altered canonical pathways in laser microdissection-enriched tumor epithelium versus stroma in the expanded cohort (n=9 cases) identified using Ingenuity Pathway Analysis.** Comparison of the most differential canonical pathways between LMD enriched tumor and stroma was determined using unsupervised analysis of 381 proteins which displayed the same direction of enrichment in all 9 patients with LIMMA adjusted p-value < 0.01 exhibiting a Log_2_ fold-change cut-off ± 1.

**Supplemental Table 26. Top altered diseases and biofunctions in laser microdissection-enriched tumor epithelium versus stroma in the expanded cohort (n=9 cases) identified using Ingenuity Pathway Analysis.** Comparison of the most differential diseases/functions between LMD enriched tumor and stroma was determined using unsupervised analysis of 381 proteins which displayed the same direction of enrichment in all 9 patients with LIMMA adjusted p-value < 0.01 exhibiting a Log_2_ fold-change cut-off ± 1.

**Supplemental Table 27. Significantly differentially expressed alterations between LMD enriched tumor epithelium versus stroma in the expanded cohort (n=9 cases) identified using Ingenuity Pathway Analysis (IPA) of known gene targets of FDA-approved anticancer drugs.** Comparison of prevalent drug targets present in LMD enriched tumor versus stroma was determined using unsupervised analysis of 381 proteins which displayed the same direction of enrichment in all 9 patients with LIMMA adjusted p-value < 0.01 exhibiting a Log_2_ fold-change cut-off ± 1. The complete list of all differentially expressed genes that are known drug targets from our proteomic-level and transcriptomic-level data was compared with a list of 150 FDA-approved anticancer drugs as of 2014 analyzed in a study by Sun et al [21].

**Supplemental Table 28. Probabilities for correlation of 343WM samples with consensus molecular subtypes using the consensusOV package in R [4].**

**Supplemental Table 29. Probabilities for correlation of samples from the expanded cohort (n=9) with consensus molecular subtypes using the consensusOV package in R [4].**

**Supplemental Table 30. Dendrogram Spearman correlations.**

**Supplemental Table 31. The top 5 ranked most and least variable proteins from genes measured in a clinical setting.** A composite list of 70 genes and/or proteins that were examined in the clinical setting for cases 343WP, 343WN, and 343WQ was generated, and variability of protein expression in cases 343WK, 343WQ, and 343WH, for which the full complement of 5 levels/case for each LMD enriched tumor, LMD enriched stroma, and whole tumor were available in our proteomic dataset. Relative standard deviation (RSD) values of protein expression were calculated between the 5 levels within each collection type within a patient’s samples and then ranked. The median rank and median RSD between patients for the top 5 most and least variable proteins within each patients’ samples are shown.

---

**Supplemental Figure 1.**
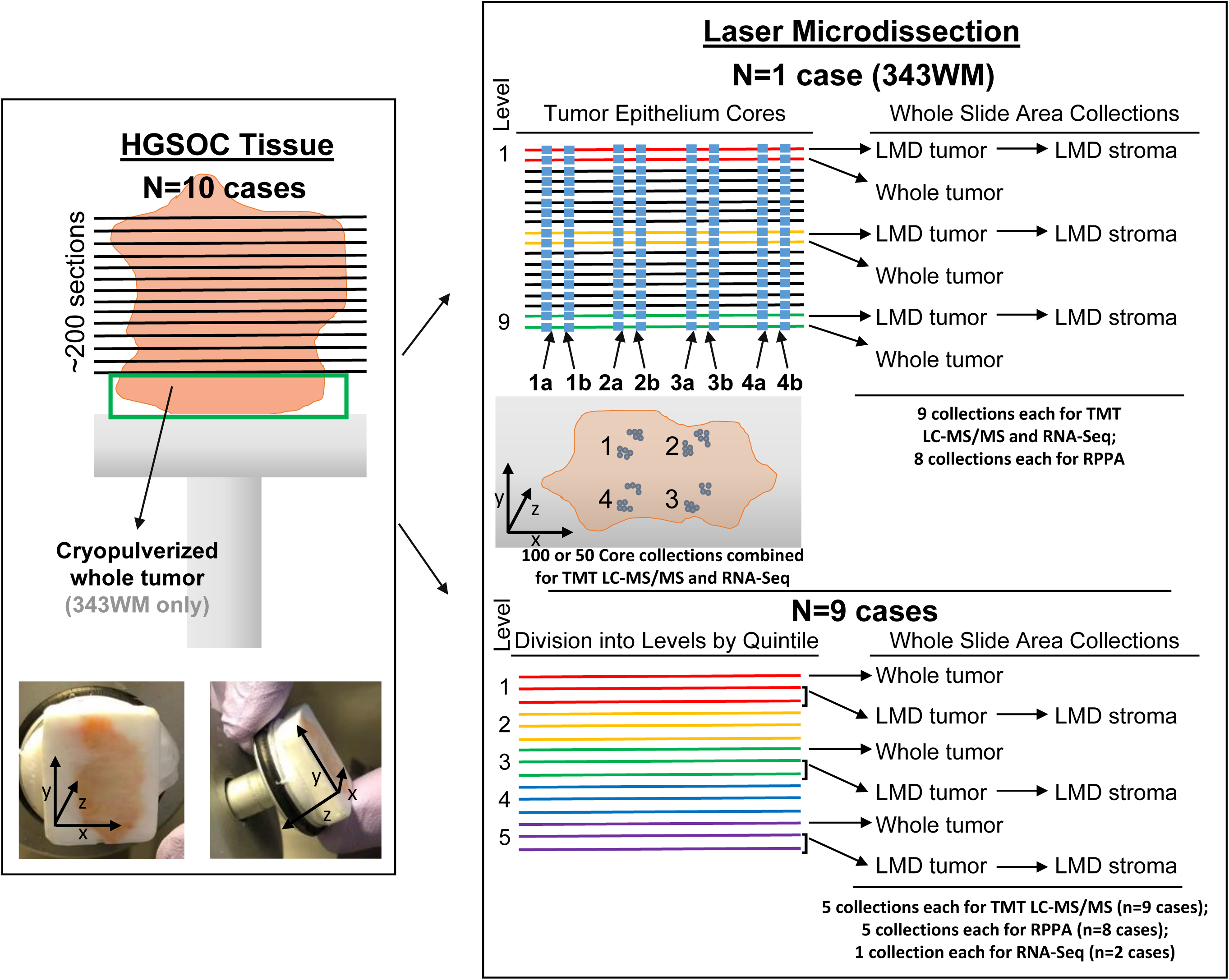
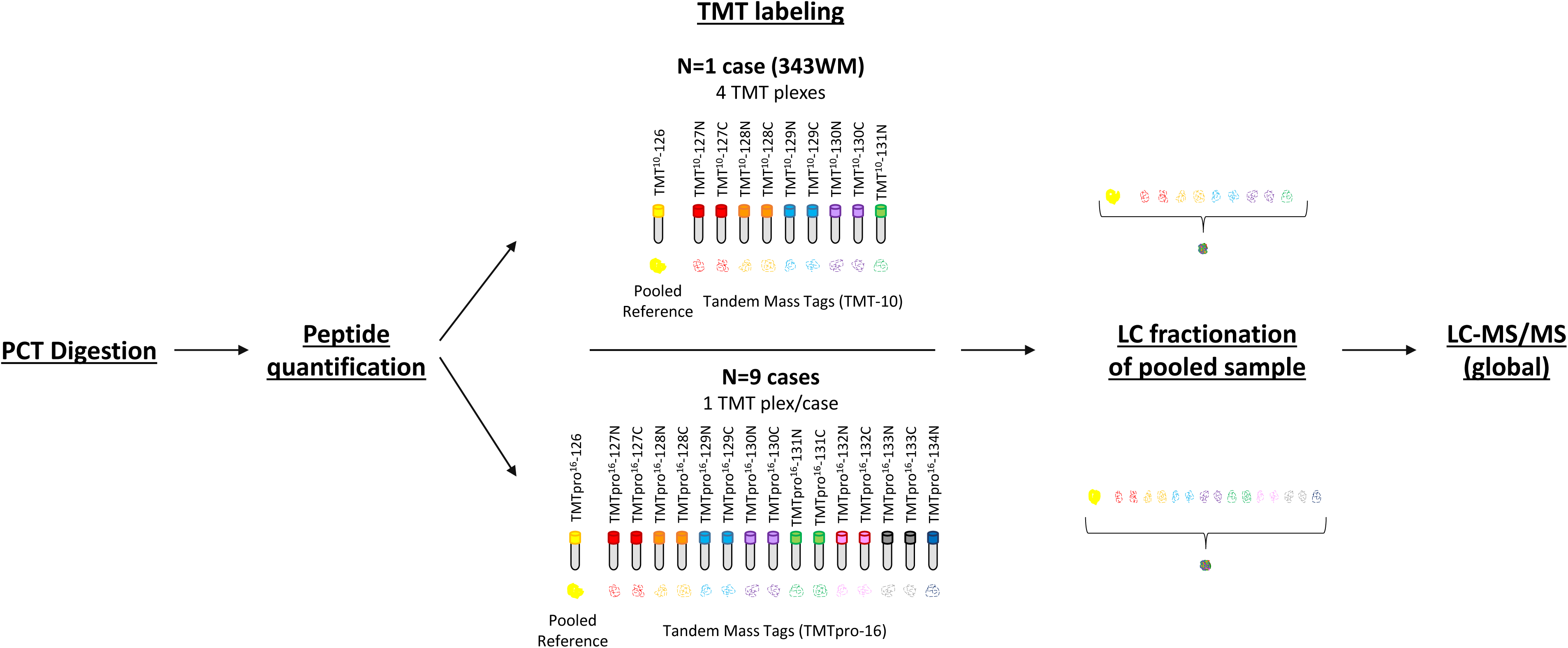
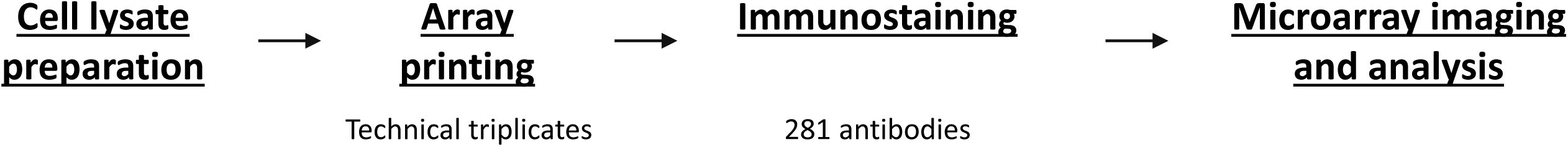
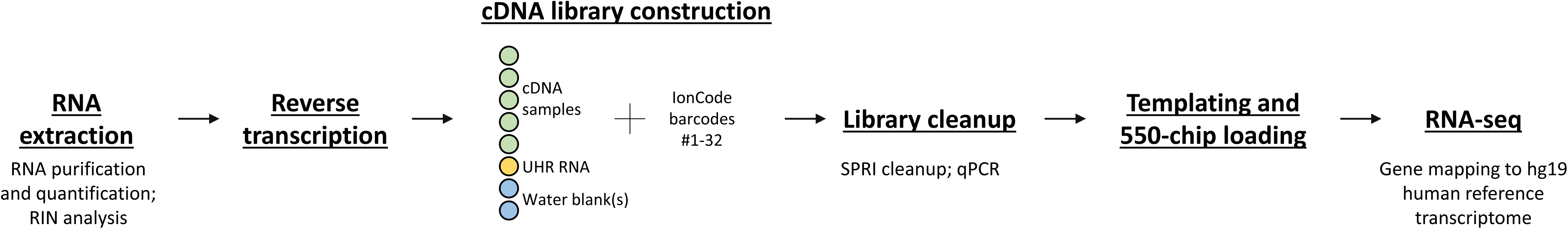
Detailed study workflow diagram. Illustration of histological tissue preparation and laser microdissection (A), isobaric tagging and high-resolution liquid chromatography-tandem mass spectrometry (TMT LC-MS/MS) analysis (B), reverse phase protein microarray (RPPA) analysis (C), and transcriptomic (RNA-seq) analysis (D). (A) Specimen blocks obtained from 10 HGSOC patients were each cut into ∼200 consecutive thin tissue sections (left panel), which were each laser microdissected for enrichment of discrete tumor epithelium, discrete stroma, or whole tumor collections (middle panel) for analysis via quantitative proteomics and transcriptomics (right panel). One case (343WM; middle panel, top) was uniquely used for laser microdissection (LMD) enrichment of four tumor epithelium cores with adjacent replicate regions from each of 100 or 50 slides evenly distributed through the depth of the specimen for MS proteomics or transcriptomics, respectively. For 343WM, additional sets of 9 slides from spatially separated levels within the specimen block were each discretely microdissected for all remaining tumor and stroma after collecting the cores by LMD, as well as a nearest neighboring whole tumor collection. The remainder of the specimen was cryopulverized in liquid nitrogen. The specimen blocks from the remaining 9 cases (middle panel, bottom) were divided into 5 levels (quintiles) of equal depth. Within each level, interlaced sections were used for LMD enrichment of discrete tumor epithelium, discrete stroma, and whole tumor collections for each downstream analytical purpose. Proteins and transcripts isolated from each of these distinct collections were analyzed by TMT LC-MS/MS (B), RPPA (C), and/or next generation sequencing (D). (B) The LMD tissue for mass spectrometry-based analysis was digested using pressure cycling technology, and digested peptides were then quantified and TMT-labeled. 10 µg/sample from case 343WM were labeled using TMT-10 plex reagents for a total of 4 TMT plexes. 5 µg/sample from the additional 9 cases were labeled using the TMTpro 16-plex reagents in patient-specific plexes. Individually labeled samples were then pooled, fractionated via basic reversed-phase liquid chromatography (bRPLC), pooled into 36 or 24 combined fractions for case 343WM or the remaining 9 cases, respectively, and pooled fractions were analyzed via LC-MS/MS. (C) LMD tissue for RPPA was collected into a tissue lysis buffer, and cell lysates were prepared for microarray printing. Samples arrayed in technical replicates (triplicates) were stained using 281 antibodies targeting native and post-translationally modified proteins. Immunostained arrays were imaged, signals were normalized relative to stained control slides, and analyzed. (D) LMD tissue for RNA-seq was collected into a lysis buffer for extraction and subsequent purification and quantification of RNA. Quality of the isolated RNA was measured. RNA samples were then reverse transcribed into cDNA and individually labeled with IonCode barcode sequences. cDNA libraries containing 5-6 LMD samples/library, plus a universal human reference RNA sample, and 1-2 control water blanks were cleaned using solid phase reversible immobilization (SPRI) to remove fragments less than 100 bp and quantified via qPCR. Library concentration was adjusted to 100 pM and used for templating and chip-loading for RNA-seq. Genes were mapped to the hg19 human reference transcriptome.

**Supplemental Figure 2.**
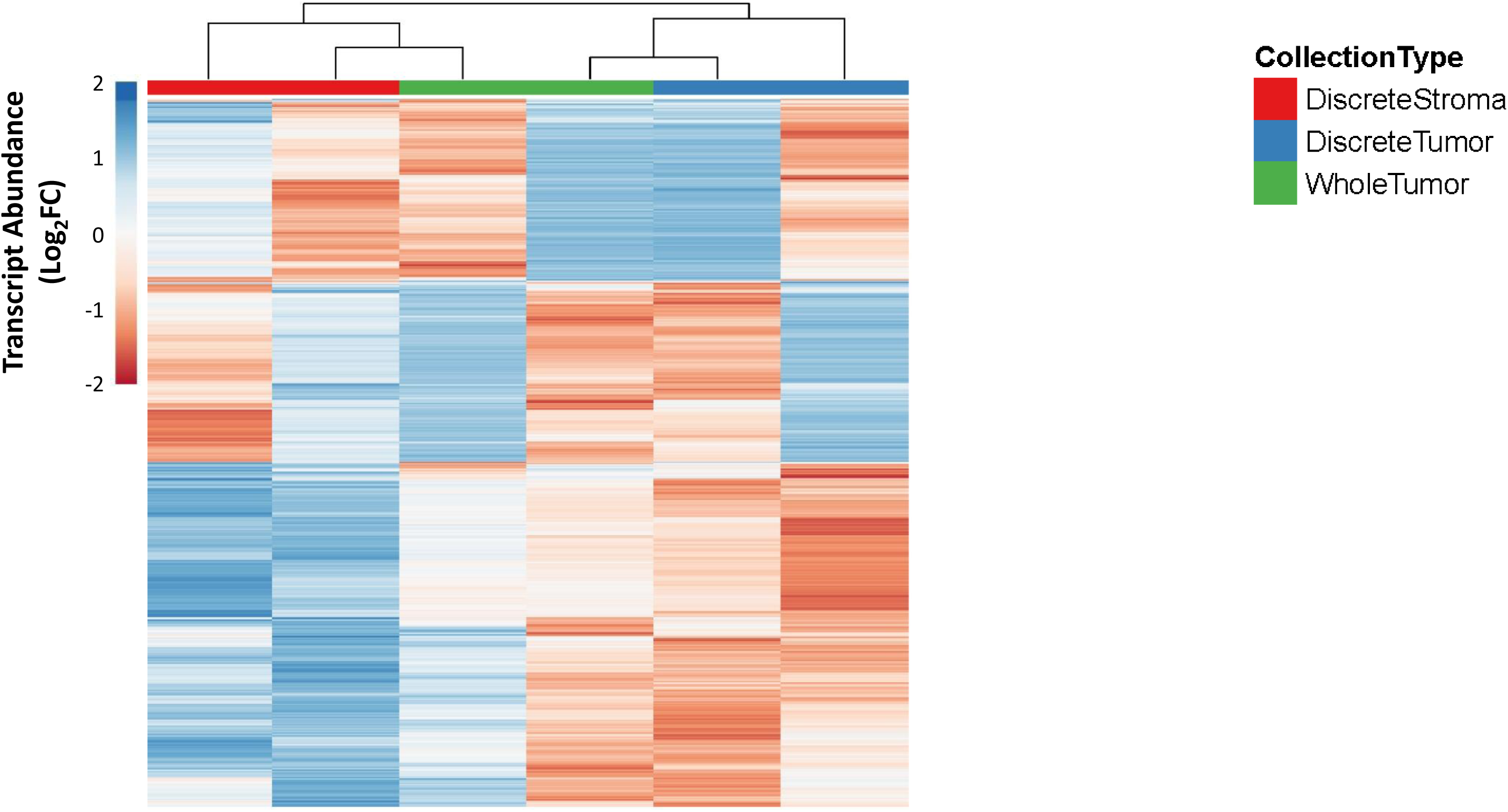
Unsupervised hierarchical cluster analysis of 874 transcripts with MAD>1 in the expanded cohort. Transcript abundances are represented across n=6 samples derived from n=2 patients in the expanded cohort consisting of: LMD enriched tumor epithelium, LMD enriched stroma, and whole tumor samples each taken from 1 level/patient.

**Supplemental Figure 3.**
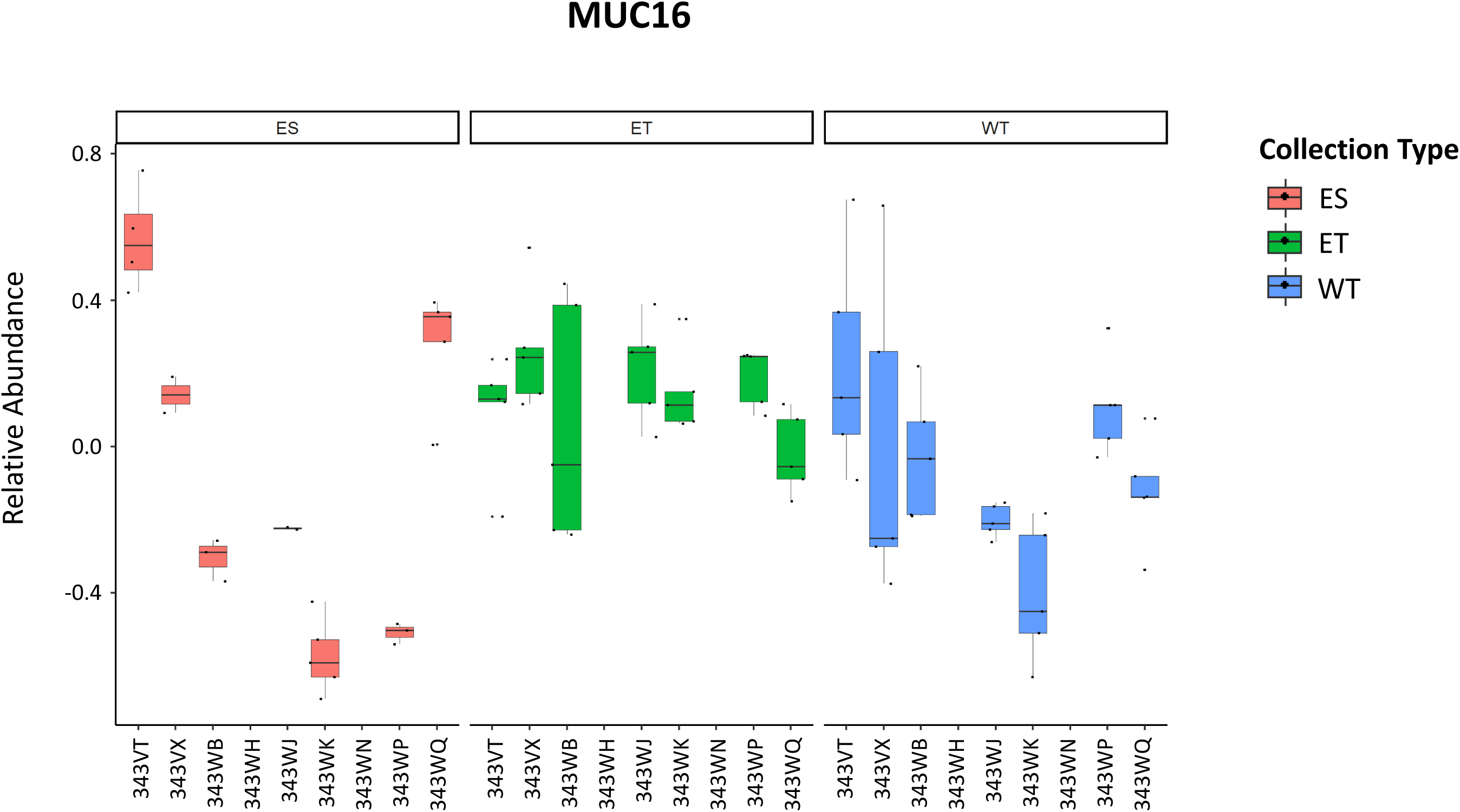
Boxplots depicting relative protein abundance for MUC16 in the expanded cohort. ES= LMD enriched stroma; ET= LMD enriched tumor; WT= whole tumor. No significant difference between ES and ET (p-value= 0.08256807).

**Supplemental Figure 4.**
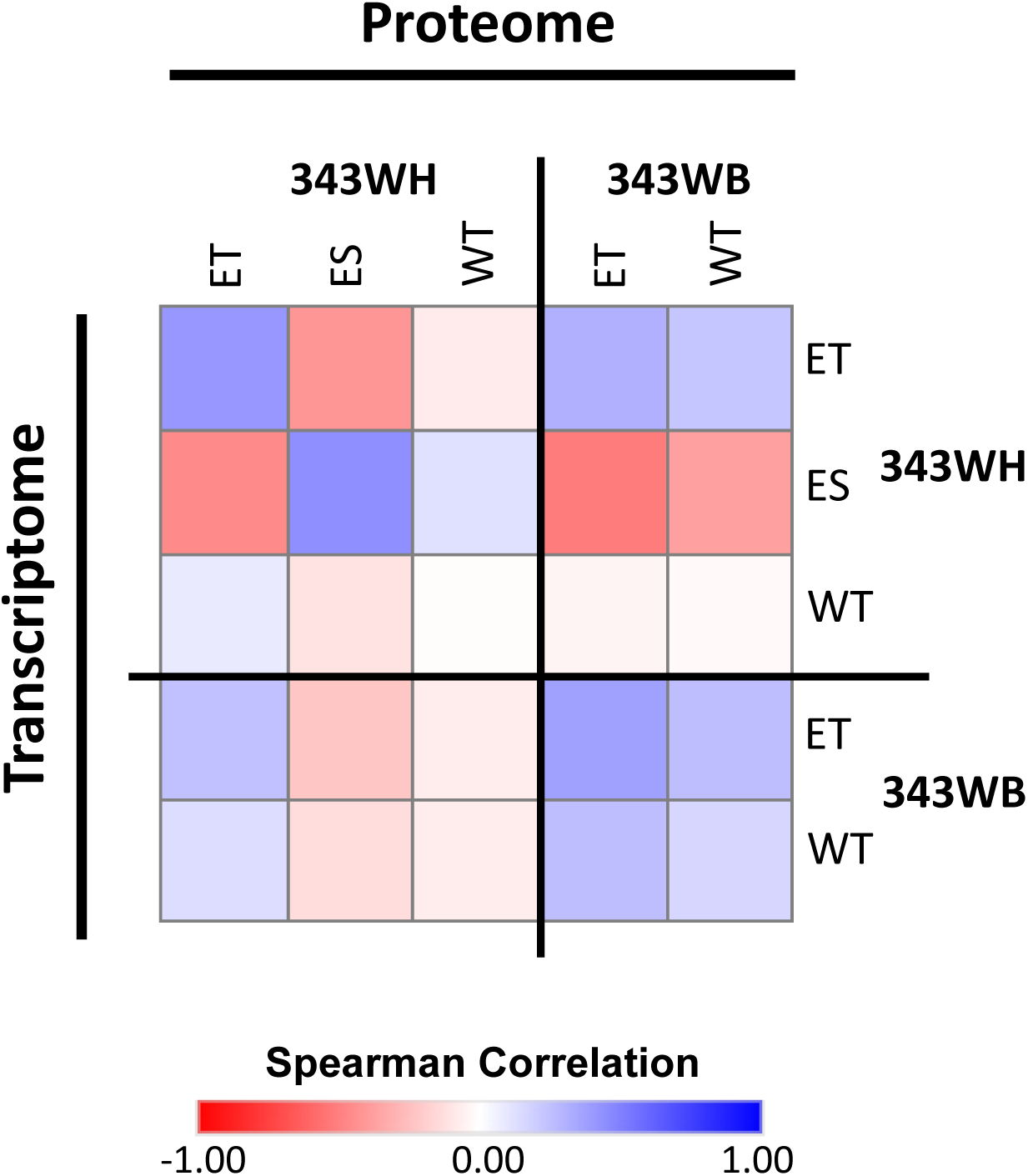
Protein-RNA Spearman Correlation Matrix for cases 343WB and 343WH. Spearman correlation analysis of 5,379 genes that were co-measured as proteins and corresponding transcripts in 343WM.

**Supplemental Figure 5.**
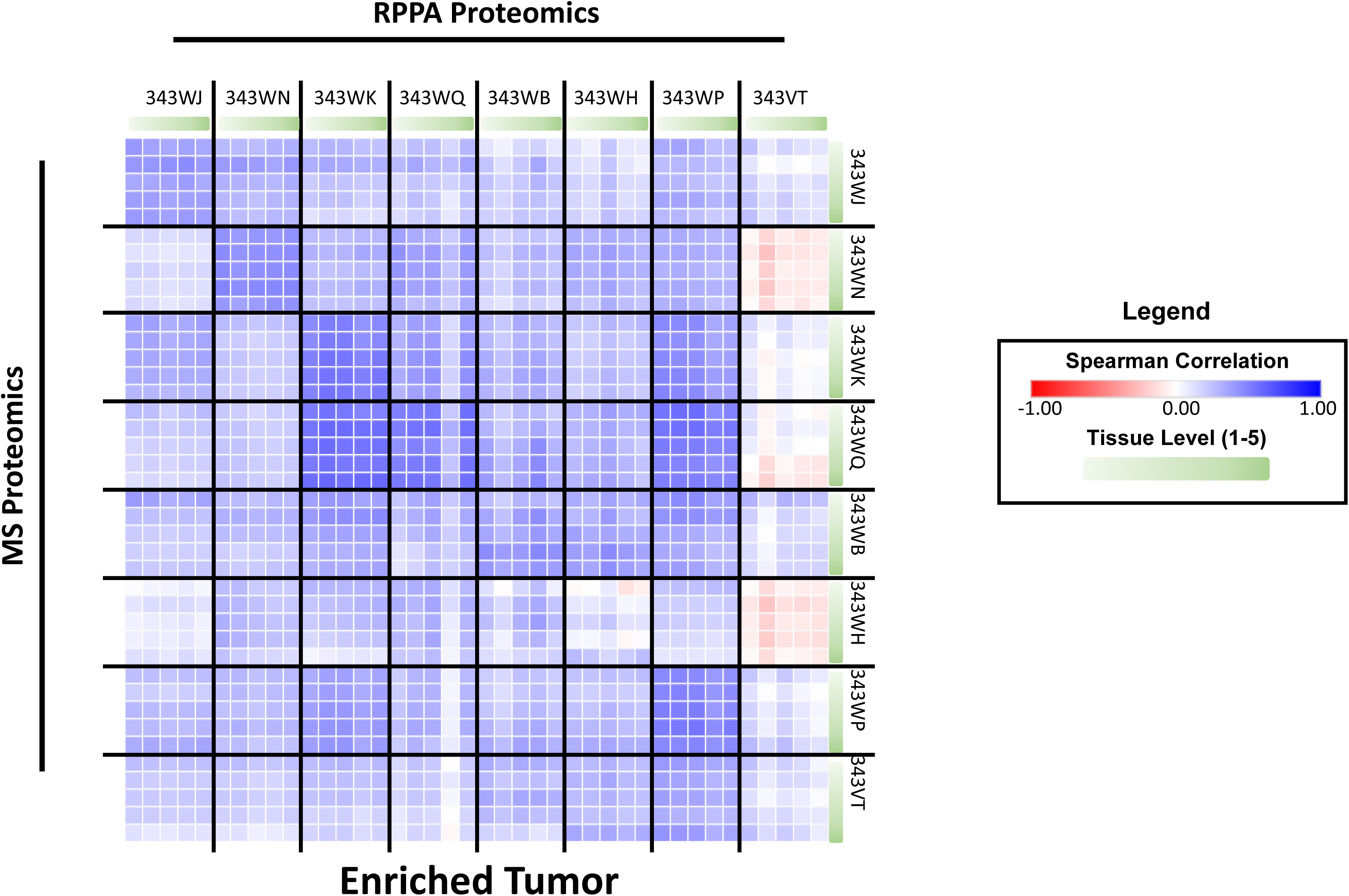
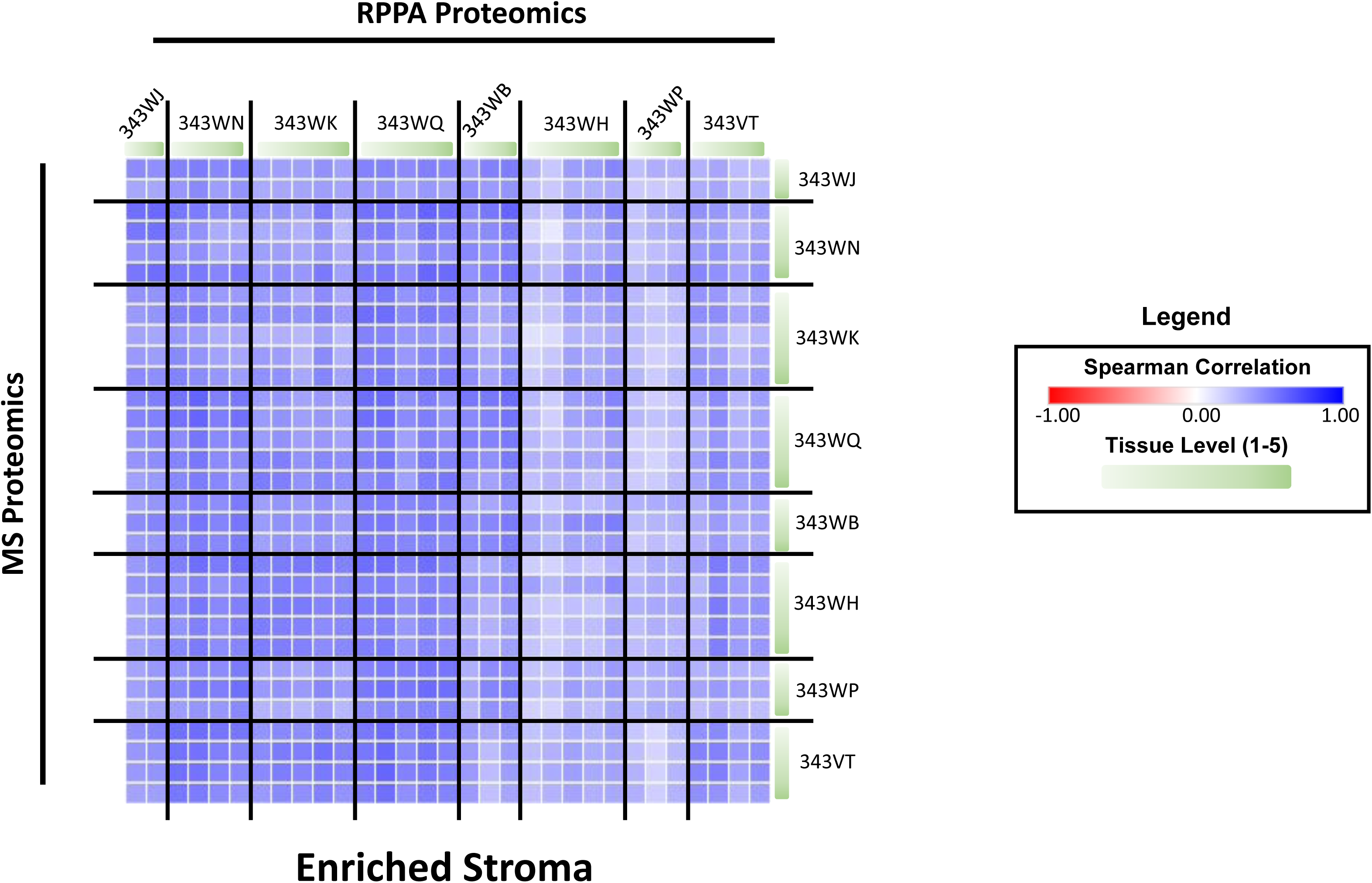
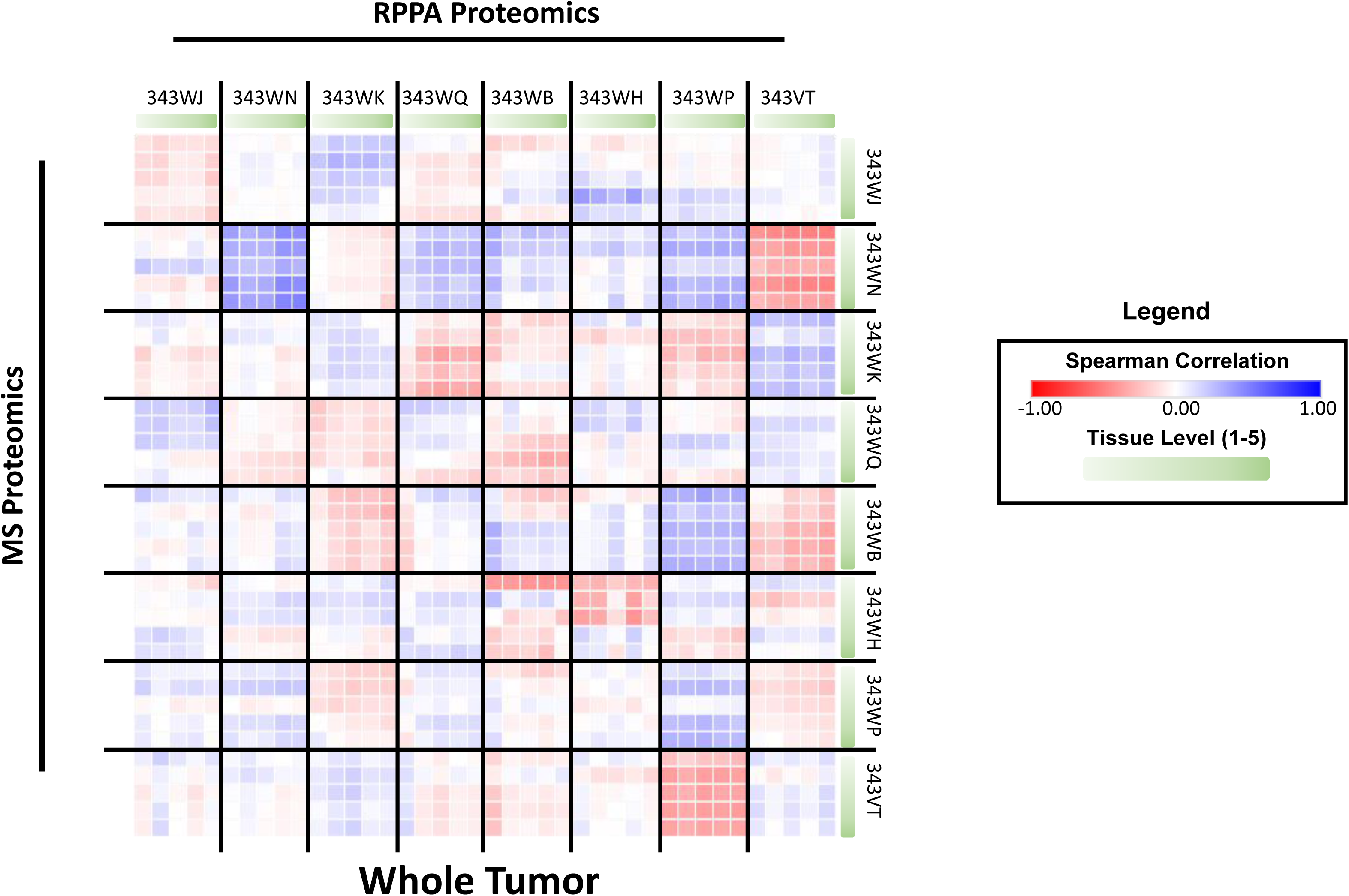
Correlation matrices of proteins co-quantified by TMT LC-MS/MS and RPPA proteomics. Spearman correlation analysis of 5,379 genes that were co-measured as proteins by TMT LC-MS/MS and RPPA for (A) LMD enriched tumor, (B) LMD enriched stroma, and (C) whole tumor.

**Supplemental Figure 6.**
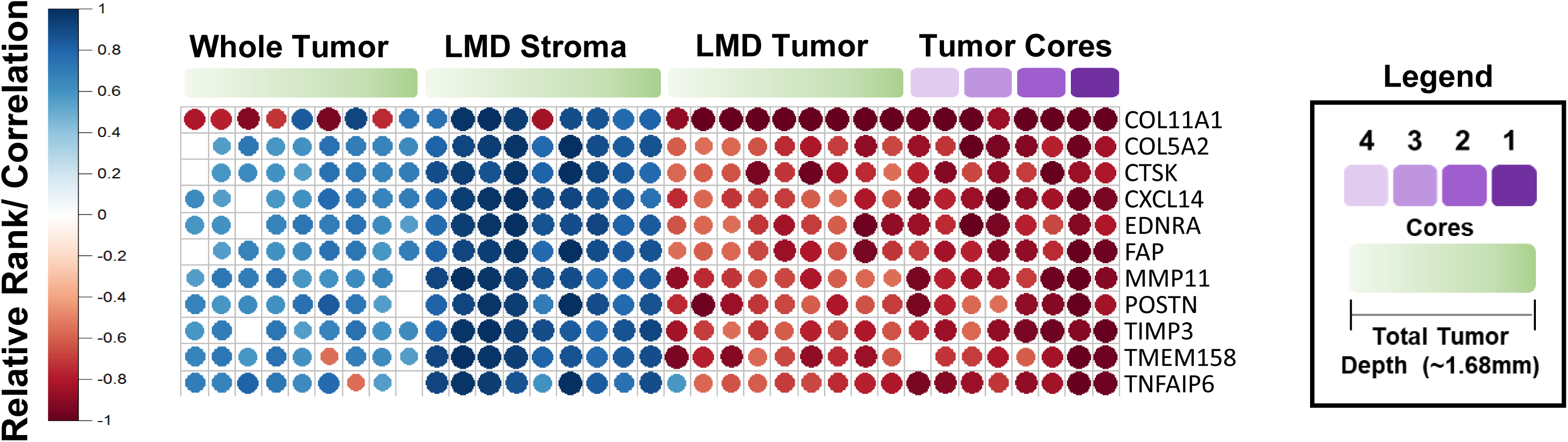
Transcript abundance of markers measured in 343WM correlating with a prognostic molecular signature of suboptimal cytoreduction in HGSOC identified by Liu et al. (2015) [24].

**Supplemental Figure 7.**
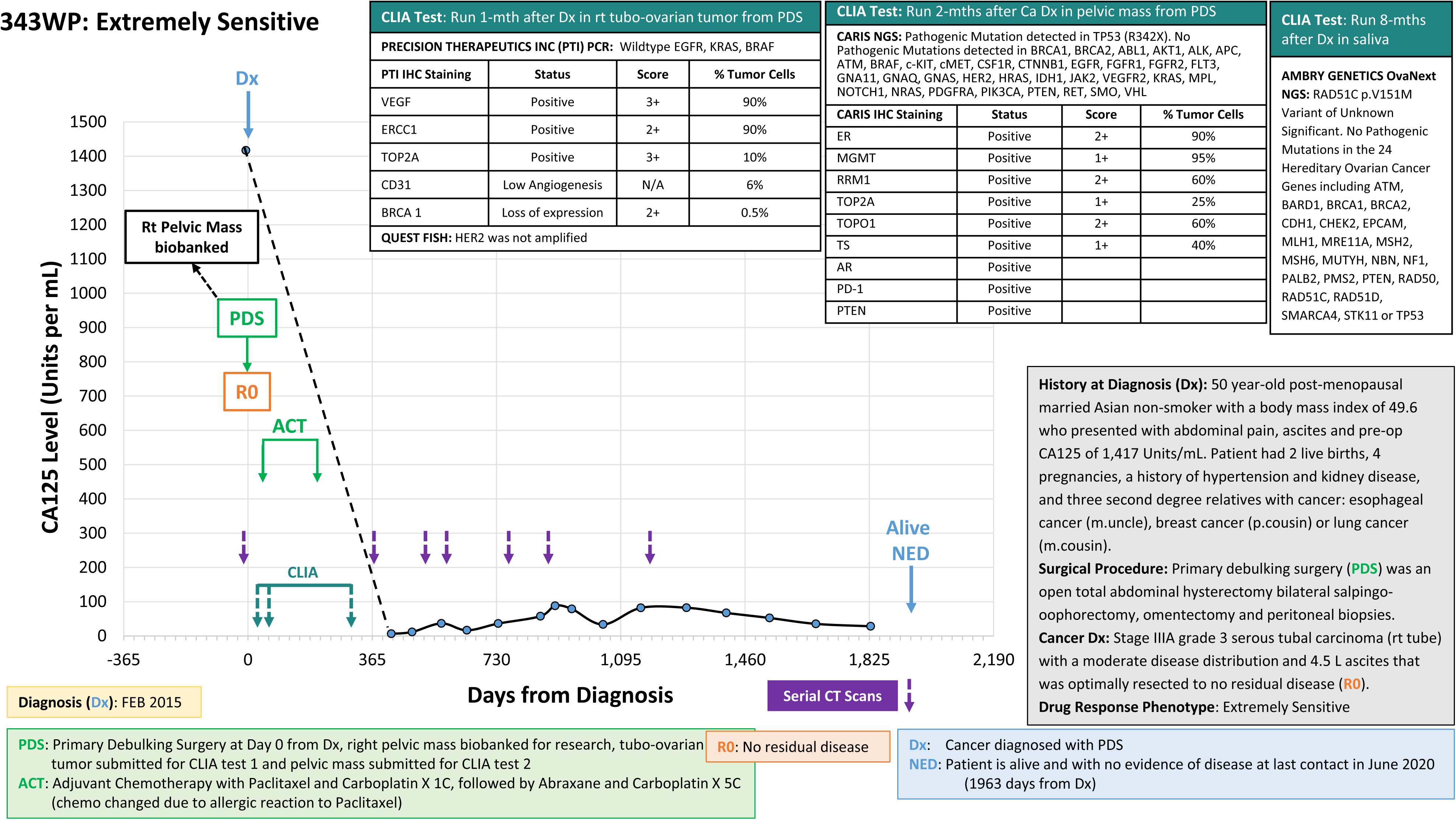
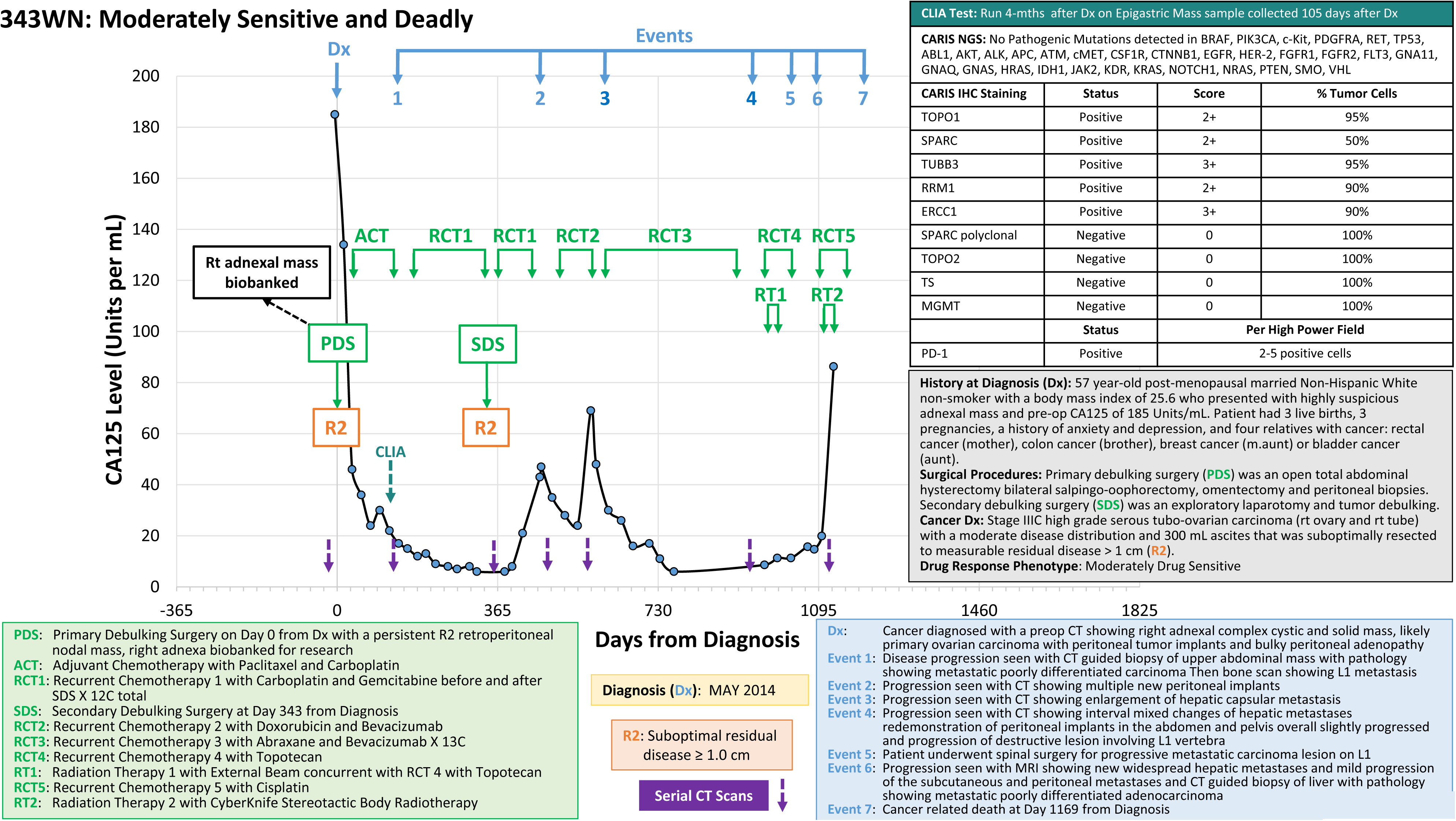
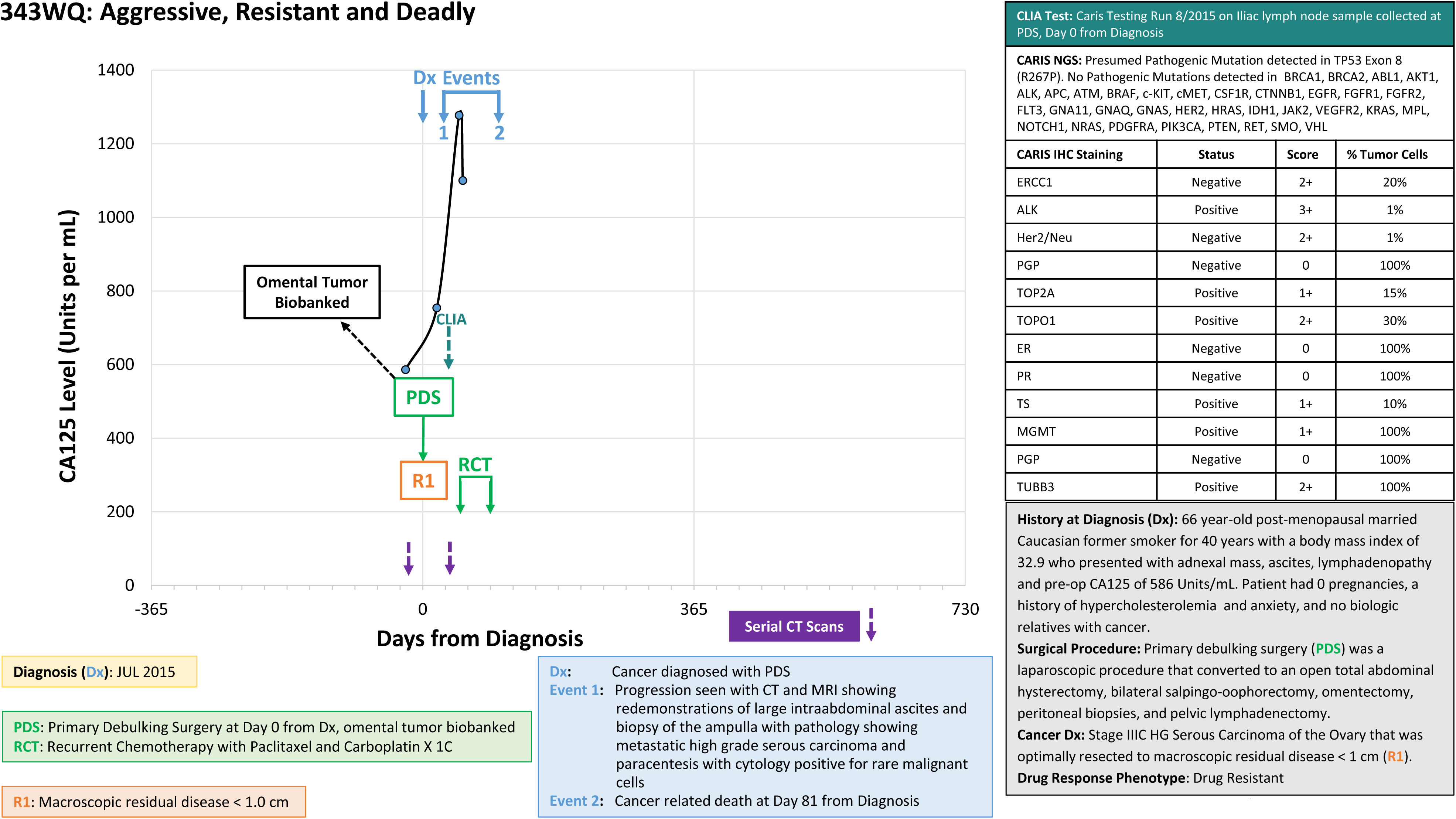
Cancer-specific clinical histories for cases 343WP, 343WN, and 343WQ. Graphs represent a timeline of each patient’s clinical history from time of diagnosis (Dx) to time of death or last follow-up and associated CA-125 levels. Brief descriptive patient histories are reported in the grey box. Dates of serial computerized tomography (CT) scans are shown as purple arrows. Dates of clinical events of diagnosis, progression recurrence, and final survival status are indicated by blue arrows and are detailed in the blue box. Dates of surgical and/or therapeutic interventions are indicated as light green arrows and are detailed in the light green box. Residual disease status following surgical debulking is indicated in orange boxes. The date acquired and tissue site for the specimens used in this study are notated in the black boxes associated with the primary debulking surgery (PDS). Dates of sample acquisition for examination of clinical marker expression and/or mutation status are indicated as dark green arrows; details of the company who analyzed the sample, the sample source (tissue or biofluid), and markers examined are listed in the CLIA test box(es) in the top right.

## References

1. Siegel, R.L., K.D. Miller, and A. Jemal, *Cancer statistics,* 2020. CA: A Cancer Journal for Clinicians, 2020. 70(1): p. 7–30.

2. Noone AM, H.N., Krapcho M, Miller D, Brest A, Yu M, Ruhl J, Tatalovich Z, Mariotto A, Lewis DR, Chen HS, Feuer EJ, Cronin KA (eds), *SEER Cancer Statistics Review,* 1975-2015. National Cancer Institute. Bethesda, MD, https://seer.cancer.gov/csr/1975_2015/, April 2018. based on November 2017 SEER data submission, posted to the SEER web site.

3. The Cancer Genome Atlas Research, N., Integrated genomic analyses of ovarian carcinoma. Nature, 2011. 474: p. 609.

4. Chen, G.M., et al., Consensus on Molecular Subtypes of High-Grade Serous Ovarian Carcinoma. Clinical Cancer Research, 2018. 24(20): p. 5037–5047.

5. Jimenez-Sanchez, A., et al., Heterogeneous Tumor-Immune Microenvironments among Differentially Growing Metastases in an Ovarian Cancer Patient. Cell, 2017. 170(5): p. 927–938 e20.

6. Schwarz, R.F., et al., Spatial and Temporal Heterogeneity in High-Grade Serous Ovarian Cancer: A Phylogenetic Analysis. PLOS Medicine, 2015. 12(2): p. e1001789.

7. Shih, A.J., et al., Identification of grade and origin specific cell populations in serous epithelial ovarian cancer by single cell RNA-seq. PloS one, 2018. 13(11): p. e0206785–e0206785.

8. Zhang, A.W., et al., Interfaces of Malignant and Immunologic Clonal Dynamics in Ovarian Cancer. Cell, 2018. 173(7): p. 1755–1769.e22.

9. Bashashati, A., et al., Distinct evolutionary trajectories of primary high-grade serous ovarian cancers revealed through spatial mutational profiling. The Journal of Pathology, 2013. 231(1): p. 21–34.

10. Verhaak, R.G.W., et al., Prognostically relevant gene signatures of high-grade serous ovarian carcinoma. The Journal of clinical investigation, 2013. 123(1): p. 517–525.

11. Yang, X., et al., FAP Promotes Immunosuppression by Cancer-Associated Fibroblasts in the Tumor Microenvironment via STAT3–CCL2 Signaling. Cancer Research, 2016. 76(14): p. 4124–4135.

12. Ghosh, S., et al., Up-regulation of stromal versican expression in advanced stage serous ovarian cancer. Gynecologic oncology, 2010. 119(1): p. 114–120.

13. Aran, D., Z. Hu, and A.J. Butte, xCell: digitally portraying the tissue cellular heterogeneity landscape. Genome biology, 2017. 18(1): p. 220–220.

14. Vergote, I.B., O.P. Børmer, and V.M. Abeler, Evaluation of serum CA 125 levels in the monitoring of ovarian cancer. American Journal of Obstetrics & Gynecology, 1987. 157(1): p. 88–92.

15. Bast, R.C., et al., A Radioimmunoassay Using a Monoclonal Antibody to Monitor the Course of Epithelial Ovarian Cancer. New England Journal of Medicine, 1983. 309(15): p. 883–887.

16. Canney, P.A., et al., Ovarian cancer antigen CA125: a prospective clinical assessment of its role as a tumour marker. British Journal of Cancer, 1984. 50(6): p. 765–769.

17. Yin, B.W.T., A. Dnistrian, and K.O. Lloyd, Ovarian cancer antigen CA125 is encoded by the MUC16 mucin gene. International Journal of Cancer, 2002. 98(5): p. 737–740.

18. O’Brien, T.J., et al., The CA 125 Gene: An Extracellular Superstructure Dominated by Repeat Sequences. Tumor Biology, 2001. 22(6): p. 348–366.

19. O’Brien, T.J., et al., The CA 125 Gene: A Newly Discovered Extension of the Glycosylated N-Terminal Domain Doubles the Size of This Extracellular Superstructure. Tumor Biology, 2002. 23(3): p. 154–169.

20. Fendrick, J.L., et al., Characterization of CA 125 Synthesized by the Human Epithelial Amnion WISH Cell Line. Tumor Biology, 1993. 14(5): p. 310–318.

21. Sun, J., et al., A systematic analysis of FDA-approved anticancer drugs. BMC systems biology, 2017. 11(Suppl 5): p. 87–87.

22. Konecny, G.E., et al., Prognostic and therapeutic relevance of molecular subtypes in high-grade serous ovarian cancer. Journal of the National Cancer Institute, 2014. 106(10): p. dju249.

23. Tucker, S.L., et al., Molecular biomarkers of residual disease after surgical debulking of high-grade serous ovarian cancer. Clinical cancer research : an official journal of the American Association for Cancer Research, 2014. 20(12): p. 3280–3288.

24. Liu, Z., et al., Suboptimal cytoreduction in ovarian carcinoma is associated with molecular pathways characteristic of increased stromal activation. Gynecologic oncology, 2015. 139(3): p. 394–400.

25. Leong, H.S., et al., Efficient molecular subtype classification of high-grade serous ovarian cancer. The Journal of Pathology, 2015. 236(3): p. 272–277.

26. Wang, C., et al., Pooled Clustering of High-Grade Serous Ovarian Cancer Gene Expression Leads to Novel Consensus Subtypes Associated with Survival and Surgical Outcomes. Clinical cancer research : an official journal of the American Association for Cancer Research, 2017. 23(15): p. 4077–4085.

27. Helland, Å., et al., *Deregulation of MYCN,* LIN28B and LET7 in a molecular subtype of aggressive high-grade serous ovarian cancers. PloS one, 2011. 6(4): p. e18064–e18064.

28. Bentink, S., et al., Angiogenic mRNA and microRNA Gene Expression Signature Predicts a Novel Subtype of Serous Ovarian Cancer. PLOS ONE, 2012. 7(2): p. e30269.

29. Torre, L.A., et al., *Ovarian cancer statistics,* 2018. CA: A Cancer Journal for Clinicians, 2018. 68(4): p. 284–296.

30. Buczak, K., et al., Spatial Tissue Proteomics Quantifies Inter-and Intratumor Heterogeneity in Hepatocellular Carcinoma (HCC). Molecular & Cellular Proteomics, 2018. 17(4): p. 810–825.

31. Zhang, Q., C. Wang, and W.A. Cliby, Cancer-associated stroma significantly contributes to the mesenchymal subtype signature of serous ovarian cancer. Gynecologic Oncology, 2019. 152(2): p. 368–374.

32. Furuya, M., Ovarian cancer stroma: pathophysiology and the roles in cancer development. Cancers, 2012. 4(3): p. 701–724.

33. Ali, M., et al., Global proteomics profiling improves drug sensitivity prediction: results from a multi-omics, pan-cancer modeling approach. Bioinformatics, 2017. 34(8): p. 1353–1362.

34. Izar, B., et al., A single-cell landscape of high-grade serous ovarian cancer. Nature Medicine, 2020.

35. Schwede, M., et al., The Impact of Stroma Admixture on Molecular Subtypes and Prognostic Gene Signatures in Serous Ovarian Cancer. Cancer Epidemiol Biomarkers Prev, 2020. 29(2): p. 509–519.

36. Bu, L., et al., Biological heterogeneity and versatility of cancer-associated fibroblasts in the tumor microenvironment. Oncogene, 2019.

37. Jia, D., et al., A COL11A1-correlated pan-cancer gene signature of activated fibroblasts for the prioritization of therapeutic targets. Cancer letters, 2016. 382(2): p. 203–214.

38. Narikiyo, M., et al., Molecular association of functioning stroma with carcinoma cells in the ovary: A preliminary study. Oncology letters, 2019. 17(3): p. 3562–3568.

39. Eckert, M.A., et al., Proteomics reveals NNMT as a master metabolic regulator of cancer-associated fibroblasts. Nature, 2019.

40. Labiche, A., et al., Stromal Compartment as a Survival Prognostic Factor in Advanced Ovarian Carcinoma. International Journal of Gynecologic Cancer, 2010. 20(1): p. 28–33-28-33.

41. Torres, D., et al., Factors that influence survival in high-grade serous ovarian cancer: A complex relationship between molecular subtype, disease dissemination, and operability. Gynecologic Oncology, 2018. 150(2): p. 227–232.

42. Hamilton, C.A., et al., The impact of disease distribution on survival in patients with stage III epithelial ovarian cancer cytoreduced to microscopic residual: a Gynecologic Oncology Group study. Gynecologic oncology, 2011. 122(3): p. 521–526.

43. Yue, H., et al., Gene signature characteristic of elevated stromal infiltration and activation is associated with increased risk of hematogenous and lymphatic metastasis in serous ovarian cancer. BMC Cancer, 2019. 19(1): p. 1266.

44. Aran, D., M. Sirota, and A.J. Butte, Systematic pan-cancer analysis of tumour purity. Nature Communications, 2015. 6: p. 8971.

45. Mao, Y., et al., Low tumor purity is associated with poor prognosis, heavy mutation burden, and intense immune phenotype in colon cancer. Cancer management and research, 2018. 10: p. 3569–3577.

46. Zhang, C., et al., Tumor Purity as an Underlying Key Factor in Glioma. Clinical Cancer Research, 2017. 23(20): p. 6279–6291.

47. Bateman, N.W., et al., Elevated AKAP12 in paclitaxel-resistant serous ovarian cancer cells is prognostic and predictive of poor survival in patients. Journal of proteome research, 2015. 14(4): p. 1900–1910.

48. Winterhoff, B.J., et al., Single cell sequencing reveals heterogeneity within ovarian cancer epithelium and cancer associated stromal cells. Gynecologic oncology, 2017. 144(3): p. 598–606.

49. Zhang, S., et al., Stroma-associated master regulators of molecular subtypes predict patient prognosis in ovarian cancer. Scientific reports, 2015. 5: p. 16066–16066.

50. Rossi, L., et al., Bevacizumab in ovarian cancer: A critical review of phase III studies. Oncotarget, 2017. 8(7): p. 12389–12405.

51. Skirnisdottir, I., T. Seidal, and H. Åkerud, The relationship of the angiogenesis regulators VEGF-A, VEGF-R1 and VEGF-R2 to p53 status and prognostic factors in epithelial ovarian carcinoma in FIGO-stages I-II. International journal of oncology, 2016. 48(3): p. 998–1006.

52. Masoumi Moghaddam, S., et al., Significance of vascular endothelial growth factor in growth and peritoneal dissemination of ovarian cancer. Cancer metastasis reviews, 2012. 31(1-2): p. 143–162.

53. Tewari, K.S., et al., Final Overall Survival of a Randomized Trial of Bevacizumab for Primary Treatment of Ovarian Cancer. Journal of Clinical Oncology. 0(0): p. JCO.19.01009.

54. Jiang, X., et al., PARP inhibitors in ovarian cancer: Sensitivity prediction and resistance mechanisms. Journal of cellular and molecular medicine, 2019. 23(4): p. 2303–2313.

55. Sun, J., et al., A systematic analysis of FDA-approved anticancer drugs. BMC Syst Biol, 2017. 11(Suppl 5): p. 87.

56. Zhao, C., et al., Prognostic values of DNA mismatch repair genes in ovarian cancer patients treated with platinum-based chemotherapy. Arch Gynecol Obstet, 2018. 297(1): p. 153–159.

57. De Geest, K., et al., Phase II clinical trial of ixabepilone in patients with recurrent or persistent platinum- and taxane-resistant ovarian or primary peritoneal cancer: a gynecologic oncology group study. J Clin Oncol, 2010. 28(1): p. 149–53.

58. Lee, S., et al., Molecular Analysis of Clinically Defined Subsets of High-Grade Serous Ovarian Cancer. Cell Rep, 2020. 31(2): p. 107502.

59. Ritchie, M.E., et al., limma powers differential expression analyses for RNA-sequencing and microarray studies. Nucleic Acids Research, 2015. 43(7): p. e47–e47.

60. Aran, D., Z. Hu, and A.J. Butte, xCell: digitally portraying the tissue cellular heterogeneity landscape. Genome Biol, 2017. 18(1): p. 220.

61. Baldelli, E., et al., *Reverse Phase Protein Microarrays*, in *Molecular Profiling: Methods and Protocols*, V. Espina, Editor. 2017, Springer New York: New York, NY. p. 149–169.

62. Baldelli, E., et al., Functional signaling pathway analysis of lung adenocarcinomas identifies novel therapeutic targets for KRAS mutant tumors. Oncotarget, 2015. 6(32): p. 32368–32379.

63. Signore, M. and K.A. Reeder, Antibody validation by Western blotting. Methods Mol Biol, 2012. 823: p. 139–55.

